# Grass wars: how native and non-indigenous *Sporobolus* battle heatwaves in salt marshes

**DOI:** 10.64898/2026.05.29.728445

**Authors:** Filippo Drigo, Pietro Antolini, Riccardo Trentin, Chiara Stefanelli, Davide Colaianni, Davide De Battisti, Chiara Frasson, Laura Airoldi, Gabriele Sales, Isabella Moro, Cristiano De Pittà

## Abstract

Heatwaves are increasing in frequency and intensity and may alter competitive interactions between native and non-indigenous plant species.
We compared the responses of the native *Sporobolus maritimus* and the non-indigenous *Sporobolus anglicus* to a simulated 5-day heatwave using an integrative approach combining morphological, physiological, biochemical, transcriptomic, and metabolomic analyses.
*S. maritimus* showed reduced survival, greater physiological damage, and no recovery, indicating high sensitivity to heat stress. Conversely, *S. anglicus* exhibited a rapid and coordinated response, limited damage to the photosynthetic apparatus, and full recovery after stress.
These results highlight the greater resilience of *S. anglicus* and suggest a potential decline of the native species in the Venetian salt marshes under increasing heat stress.

**Graphical Abstract:** 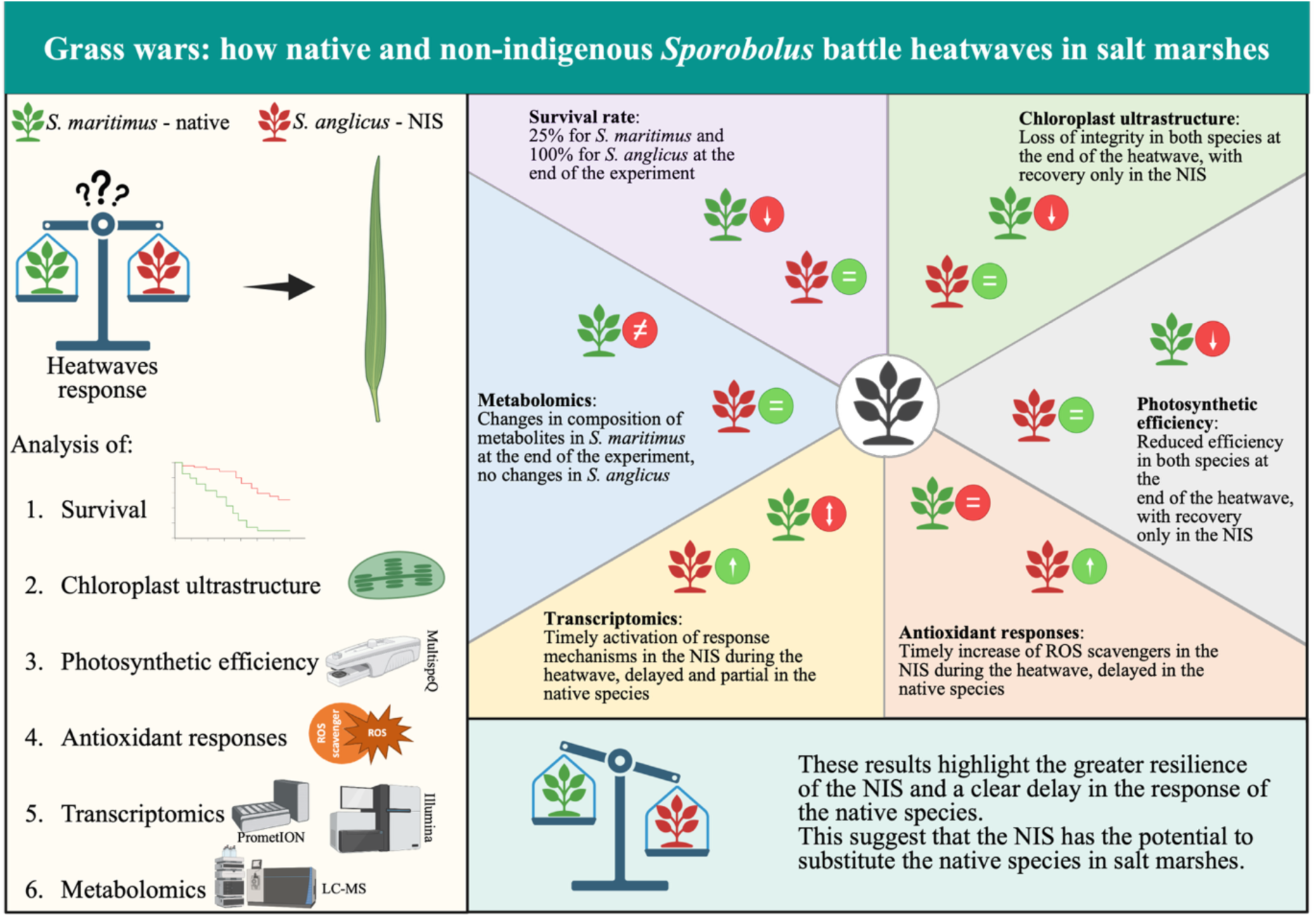

We investigated the effects of heatwaves on a native and NIS *Sporobolus* species from the Venice Lagoon using an integrative, multi-level approach during a simulated 5-day heatwave and recovery phase. The native species showed reduced survival, pronounced physiological damage, and no recovery, indicating high sensitivity to heat stress. Conversely, the NIS exhibited a rapid response, limited impairment of the photosynthetic apparatus, and full recovery. These findings indicate greater resilience of the NIS and suggest a potential decline of the native species in the Venice Lagoon under increasing heat stress. Image created with BioRender (www.biorender.com).

## 1. Introduction

*Sporobolus maritimus* (Curtis) P.M.Peterson & Saarela (autochthonous) and *Sporobolus anglicus* (C.E.Hubb.) P.M.Peterson & Saarela (allochthonous) coexist in the Venice Lagoon since the early beginning of the 21^st^ century, when first discoveries of non-native *Sporobolus* species were reported (Scarton *et al*., 2003; Ghirelli *et al*., 2004). These species play an important ecological role in sediment dynamics of salt marshes, where they are considered “ecological engineers” due to robust shoots, rhizomes, and root system (Ainouche *et al*., 2012). However, the two species compete for the same ecological niche (Wong *et al*., 2018). It was observed that *S. anglicus* rapidly spread across salt marshes in continental Europe, UK, and worldwide (Baumel *et al.,* 2001; Morgan & Systma, 2010; Maebara *et al*., 2020), with a parallel regression of the local species (Goss-Custard & Moser, 1988). Specifically, *S. anglicus* has proliferated in various areas of the Venice Lagoon, including the northern, central, and southern areas. Recent monitoring activities conducted by our research group (manuscript in preparation) and others (Wong *et al*., 2018) indicate that this species now appears to be more prevalent than the native species. This might represent a threat for native population of *S. maritimus*.

Transitional Water Systems (TWS) as the Venice Lagoon are facing a significant risk of local biodiversity loss. The major threats are invasion of Non-Indigenous Species (NIS), and anthropogenic activities inducing habitat loss, pollution, overexploitation of biological resources, and climate change (Hobday *et al*., 2016; Barriopedro *et al*., 2023). As TWS ecosystems possess unique gradients of environmental conditions, to which autochthonous species are adapted, the above-mentioned threats are of increasing concern (Cuthbert *et al*., 2021). The Venice Lagoon is one of the biggest transitional ecosystems in the Mediterranean Sea, and is affected by extreme climatic events, which are reported more frequently in the last decades (Ferrarin *et al*., 2024). Among these events, heatwaves are one of the main concerns in relation to the ongoing climate change (Lhotka *et al*., 2018). Perkins *et al*. (2012) reported an increase in frequency, intensity, and duration of heatwaves in the interval 1950-2011, which is likely caused by greenhouse gases emission (Garrabou *et al*., 2022; Barriopedro *et al*., 2023). Furthermore, climate models predict this trend will continue for at least the next 100 years, with temperature extremes becoming even more common, severe, and prolonged (Perkins & Alexander, 2013). Moreover, global climate change will make heatwaves longer, more intense, more widespread, and more frequent (Barriopedro *et al*., 2023). Some projections suggest up to three or four heatwaves per summer, with severe events becoming a regular occurrence (Lhotka *et al*., 2018). Particularly, the Mediterranean Sea is warming more than any other area of the world, with an average warming rate between 1982 and 2019 of 0.38 °C per decade, compared to a global average of 0.11 °C per decade (Garrabou *et al*., 2022). This makes the Mediterranean Sea more susceptible to heatwaves (Barriopedro *et al*., 2023). More specifically, the Venice Lagoon has been affected by several heatwaves from the beginning of the 21^st^ century (Ferrari *et al*., 2024). This might cause severe heat stress on local species, including plants (Barriopedro *et al*., 2023).

Within this context, we hypothesize that the NIS *S. anglicus* is more tolerant to heatwave-related stress than the native species *S. maritimus*. This hypothesis is supported by the observed expansion of *S. anglicus* within the lagoon (Wong *et al*., 2018) and may have an evolutionary explanation. *S. anglicus* arose from chromosome doubling of the hybrid *S. x townsendii*, which originated in the late 19th century from the hybridization of *S. maritimus* with *S. alterniflorus* (Ainouche *et al*., 2004; Ainouche *et al*., 2012). Chromosome doubling is known to enhance physiological tolerance and stress resilience in plants, in line with the theory of hybrid vigor (Chen, 2010; Botet & Keurentjes, 2020), as exemplified by the higher heat-stress resistance observed in *Triticum aestivum* hybrids (Al-Ashkar *et al*., 2020).

To test our hypothesis, we have simulated a heatwave in laboratory-controlled conditions and analyzed the response of *S. maritimus* and *S. anglicus* at the morphological, physiological, transcriptomic, and metabolic level. While morphological traits and physiological parameters have traditionally been used to assess plant stress responses, emerging omics approaches, such as transcriptomics and metabolomics, provide deeper insights into the biological processes affected by environmental stress. These methods help identify key mechanisms underlying phenotypic plasticity, acclimation, and long-term adaptation (Gleason & Burton, 2015; Oomen & Hutchings, 2017; Selechnik *et al*., 2019). Transcriptomic analyses reveal early changes in gene expression (Lee & Yeom, 2023), while untargeted metabolomics provides a comprehensive view of plant metabolism (Bittremieux *et al*., 2024). The multi-disciplinary approach used in this study has effectively elucidated heat response mechanisms in both species under investigation.

## 2. Materials and Methods

### 2.1 Plants collection

Shoots of *S. maritimus* and *S. anglicus* were collected in May of 2024 from a salt marsh in the Venice Lagoon (45.356948° N, 12.229714° E). After transfer to the laboratory, shoots were cleaned from the salt marsh soil and placed in pots with perlite. Afterwards, we prepared a total of 4 pots per species, with 12 shoots per pot.

### 2.2 Plants identification

To distinguish the two species, both a molecular marker and nuclear genome size were analyzed. Genomic DNA (gDNA) was extracted from a single shoot per pot using the DNeasy® Plant Pro Kit (Qiagen, Germany), following the manufacturer’s protocol with minor modifications. For molecular identification, a two-locus approach was employed, using one nuclear marker (*Internal Transcribed Spacer, ITS*) and one plastid marker (intergenic spacer region *trnL-trnF*), as these provide a reliable tool to distinguish the two species (Muhammad & Ki, 2022). Primer sequences are reported in Table S1.

Nuclear genome size was estimated by flow cytometry from ∼100 mg of fresh leaf tissue from the same shoot. Six sample groups were prepared: pool M (leaves from each *S. maritimus* pot), pool A (leaves from each *S. anglicus* pot), and four mixed pairs (one leaf per species from different pots). Tissue was chopped (∼5 chops/second) in 1 mL cold nuclear extraction buffer (Baumel *et al*., 2003), filtered (50 µm), treated with RNase A (50 µg/mL, for 1 h, at Room Temperature/RT), stained with propidium iodide (50 µg/mL, for 15 min, on ice, in darkness), and analyzed using a Cytoflex Flow Cytometer (Beckman Coulter) using the CytExpert software (Beckman Coulter). PI fluorescence was measured at 488 nm (610/20 filter). We performed a first gate on PI *vs*. SS (Side Scatter) and then we considered the expression of PI on a histogram PI *vs*. count. The sampling lasted till 10^4^ nuclei/events per sample were recorded. All sampled individuals were validated with this integrative approach (Figure S1 and Table S2).

### 2.3 Conditions of the simulated heatwave

After cleaning from lagoon soil, pots were transferred in a growth chamber (POL-EKO, Poland). For the whole experiment, the light:dark cycle was of 16:8 hours, with light intensity of 250 µmol/(m^2^*s) of photons, and plants were irrigated daily with 100 mL per pot of ¼ Hoagland’s solution (Hoagland & Arnon, 1950). To avoid drought stress during any experimental condition, pots were kept on flowerpot dishes to keep the perlite damp. Before the heatwave simulation, plants were acclimated for 10 days at environmental conditions (25 °C during the day and 23 °C during the night). Heatwave conditions were simulated by exposing the plants to 38 °C during the day and 34 °C during the night for 5 consecutive days. These conditions were defined based on the heatwave criteria provided by Perkins & Alexander (2013), using temperature records from the Venice Lagoon (Città di Venezia, 2023) and previous experimental studies on *Sporobolus* species (Duarte *et al*., 2016). Following the 5-day exposure, a recovery phase was simulated by gradually returning temperatures to initial conditions in two steps: 3 days at 30 °C (day) and 28 °C (night), followed by 3 days under acclimatation conditions (Figure 1).

**Figure 1.**
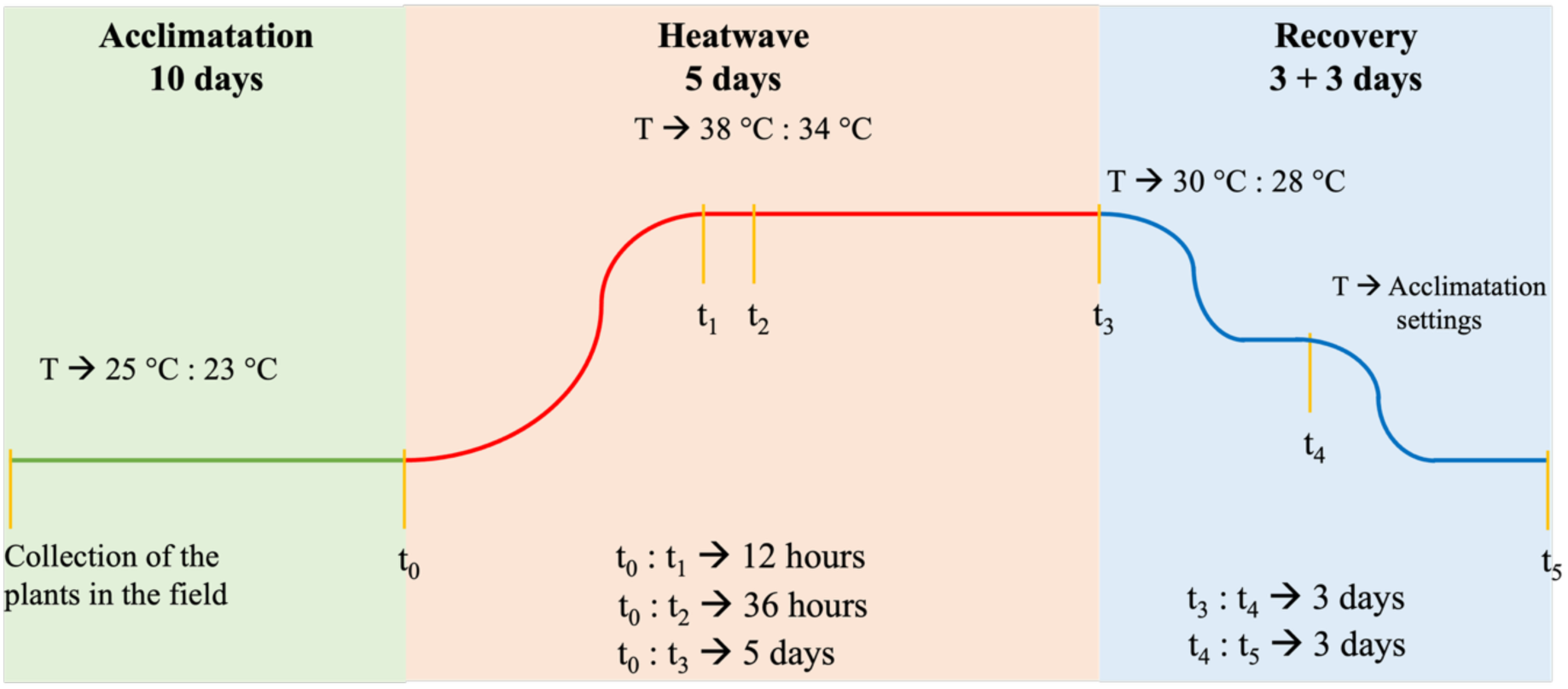
Experimental design of a laboratory-controlled simulated heatwave applied to *S. maritimus* and *S. anglicus*. The schematic illustrates the temperature regimes during different experimental phases: Acclimatation (25 °C by day and 23 °C by night), Heatwave (38 °C by day and 34 °C by night), and Recovery (30 °C by day and 28 °C by night for three days and 25 °C by day and 23 °C by night for the last three days), their durations, and the specific time points designated for sample collection. See Figure S2 for the temperature (°C) variation detected by the cultivation chamber during the experiment.

### 2.4 Photosynthetic efficiency

Photosynthetic efficiency was evaluated considering: (1) F_v_’/F_m_’, the maximum quantum yield of photosystem II (PSII); (2) ϕII, the effective photochemical efficiency of photosystem II; (3) ϕNPQ, non-photochemical quenching.

To assess photosynthetic efficiency, chlorophyll fluorescence was measured using the MultispeQ V2.0 device (PhotosynQ, USA). The “Rides 2.0” protocol was employed for the analysis, and a custom light guide mask, constructed according to instructions from the PhotosynQ website, was applied. This analysis was performed on four biological replicates.

### 2.5 Photosynthetic pigment content

#### 2.5.1 Chlorophylls and total carotenoids

Leaf samples were collected and stored at -20 °C until extraction. Approximately 5 mg of tissue was homogenized with a metal bead using a TissueLyser LT (Qiagen, Germany; 35 Hz, 30 s), repeating the cycle until complete disruption. Pigments were extracted in 2 mL of 90% HPLC-grade acetone, followed by an additional disruption step, and samples were incubated overnight at 4 °C in the dark. After centrifugation (13’500 × g, 5 min, 4 °C), supernatants were collected and absorbance was measured at 480, 510, 630, 647, and 664 nm using a UV-1200 UV/VIS spectrophotometer (AOE Instruments Co., China). Chlorophyll *a* (chl *a*), chlorophyll *b* (chl *b*), and carotenoids concentrations were calculated according to Bai *et al*. (2011). Analyses were performed on four biological replicates.

#### 2.5.2 High-Performance Liquid Chromatography (HPLC)

Acetone extracts from leaf samples (used for chlorophylls and carotenoids analyses) were further analyzed by High-Performance Liquid Chromatography (HPLC) using an Agilent 1100 system (Agilent Technologies, USA) equipped with a UV-VIS detector and a C_18_ column (5 μm particle size; 25 × 0.4 cm; 250/4 RP 18) (Merck, Germany). Aliquots (250 μL) were injected and separated using two mobile phases: A (86.7% acetonitrile, 9.6% methanol and 3.6% Tris-HCl, pH 7.8) and B (80% methanol and 20% hexane). The gradient was as follows: 100% A from 0-12.5 min, 100% B from 12.5-20.5 min, and re-equilibration with 100% A from 20.5-23 min. All steps were performed with a flow rate of 2 mL/min. Compounds were identified based on retention time and absorption spectra (Färber & Jahns, 1998). Analyses were performed on three biological replicates.

### 2.6 Transmission Electron Microscopy (TEM)

The photosynthetic membrane ultrastructure was evaluated through the observation of leaf samples at the TEM. Small pieces of leaves (approximately 0.5 × 0.5 cm squares) were collected and transferred in a 1.5 mL tube containing a fixing solution of 6% glutaraldehyde in 0.1 M sodium cacodylate buffer (pH 6.9). After fixation, samples were washed 2-4 times with 0.1 M sodium cacodylate buffer (pH 6.9) to remove the excess glutaraldehyde. Subsequently, samples were postfixed in 1% OsO_4_ for 2 h and dehydrated in a graded series of ethyl alcohol and propylene oxide and embedded in araldite according to Moro *et al*. (2020). For TEM observations, ultrathin (600 nm) sections were cut using a Ultracut Reichert-Jung ultramicrotome (Leica, Germany). These sections were then post-stained with uranyl acetate and lead citrate. The prepared samples were examined and photographed using a Tecnai G2 microscope (FEI Company, USA) operating at 100 kV.

### 2.7 Oxidative assays

#### 2.7.1 Preparation of leaf material

For the oxidative assays, approximately 50 mg of leaf samples were chopped with sterile scalpels, flash-frozen in liquid nitrogen and then stored at -80 °C until the analysis. The samples (∼10 mg for CCA and ∼50 mg for the other assays) were disrupted with metal beads using a TissueLyser LT (Qiagen, Germany) at a frequency of 35 Hz for 30 seconds, repeating the procedure until the complete fragmentation of the leaf tissue. After this step, the extraction buffer, specific for each assay, was added. Total soluble protein content was determined using the Bradford assay (Bradford, 1976), adapted to a 96-well plate format. Absorbances at 595 nm (A_595_) was measured using a Spark microplate reader (Tecan, Switzerland). Protein concentrations (µg/mL) were calculated from a calibration curve generated with bovine serum albumin (BSA).

#### 2.7.2 Ascorbate Peroxidase (APx), Catalase (CAT), and Guaiacol Peroxidaes (GPx)

The activity of APx, CAT, and GPx was determined following Hessini *et al*. (2009) for extraction and reaction buffer preparation and Duarte *et al*. (2014; 2016) for absorbance measurements, which were carried out using a UV-1200 UV/VIS spectrophotometer (AOE Instruments Co., China). Enzyme activities were expressed as units (U) per µg of extracted proteins (U/µg of proteins). All analyses were performed on four biological replicates.

#### 2.7.3 Superoxide Dismutase (SOD)

The activity of SOD was determined using the “Superoxide Dismutase (SOD) Colorimetric Activity Kit” (Thermo Fisher Scientific, USA), following the manufacturer’s guidelines on a 96-well plate, using a Spark microplate reader (Tecan, Switzerland). Results were expressed as enzyme activity units (U) per mg of extracted proteins (U/mg of proteins). The analysis was performed on four biological replicates.

#### 2.7.4 Copper chelating activity (CCA)

CCA was determined by adding 2 mL of 99% ethanol to the disrupted tissue and incubating overnight at 20 °C under agitation. Subsequent reaction steps were performed according to Trentin *et al*. (2020) using a 96-well plate format. Absorbance was measured at 632 nm with a Spark microplate reader (Tecan, Switzerland). Results were expressed as mg of Trolox equivalents (TE) per mg of fresh weight (FW) (mg TE/mg of FW), based on a calibration curve prepared with Trolox standard solutions. All analysis were performed on four biological replicates.

### 2.8 Total RNA extraction

Approximately 50 mg of leaf samples were chopped with sterile scalpels, flash-frozen in liquid nitrogen and then stored at -80 °C. The samples were disrupted with a metal bead using a TissueLyser LT (Qiagen, Germany) at a frequency of 35 Hz for 30 seconds, repeating the procedure until the complete fragmentation of the leaf. Total RNA was extracted from each sample by using the TripleXtractor reagent (GRiSP, Portugal) and the miRNeasy Mini Kit (Qiagen, Germany) according to manufacturers’ instructions. RNA concentration and purity were measured using the NanoDrop 2000c spectrophotometer (Thermo Fisher Scientific, USA), and RNA integrity was assessed by electrophoresis using the Agilent 4150 TapeStation (Agilent Technologies, USA).

### 2.9 RNA sequencing (RNA-seq) and processing

#### 2.9.1 Long-read sequencing and processing

RNA-seq analysis for the *de novo* transcriptome assembly was performed by IGATech (Udine, Italy) on leaf samples. Approximately 300 ng of total RNA per sample was used for library preparation following the PCR-cDNA Barcoding Kit v14 SQK-PCB111.24 (Oxford Nanopore, UK) protocol. Sequencing was carried out on a PromethION 2 Solo platform (Oxford Nanopore, UK) using standard settings and MinKnow software (v24.02.16).

Basecalling was performed with Guppy (v6.3.9) in high-accuracy mode, applying a minimum quality score threshold of 9. Adapter trimming was conducted using using Porechop (v0.2.4) (Wick *et al*., 2017) with default settings, except for a minimum split read size of 700 bp, and a minimum read length (-m) of 100 bp. Raw sequencing data are available in the NCBI BioProject database under accession number PRJNA1269774.

#### 2.9.2 Short-read sequencing and processing

The RNA-seq analysis of leaf samples collected during the simulated heatwave experiment was performed by Novogene (Munich, Germany) on three biological replicates per condition. Libraries were prepared using the Novogene NGS Stranded RNA Library Prep Set kit. Library quality and quantity were assessed using Qubit and real-time PCR, while fragment size distribution was evaluated with a Bioanalyzer. Libraries were then pooled and sequenced in paired-end mode (150 bp) on a NovaSeq X Plus platform (Illumina, USA).

Raw reads were pre-processed using Cutadapt (v4.9) (Martin, 2011) in paired-end mode to remove Illumina adapters from both the 5’ and 3’ ends. Sequencing data are available in the NCBI BioProject database under accession number PRJNA1269774.

### 2.10 *De novo* transcriptome assembly

Long reads were used to generate a *de novo* transcriptome assembly using RNA-Bloom2 (v1.3.1), which employs strobemers for reference-free assembly (Nip *et al*., 2023). To improve assembly accuracy, the option to incorporate short reads was enabled, using Illumina data from the simulated heatwave experiment. These reads were combined into two comprehensive files (R1 and R2) for each species (*S. anglicus* and *S. maritimus*). To minimize potential contamination from non-coding RNA fragments in downstream analyses, short transcripts generated by RNA-Bloom2 were removed from the final assemblies. Transcript redundancy was further reduced using CD-HIT (v4.8.1) (Li & Godzik, 2006) with an identity threshold of 80% (-c 0.8), word size 5 (-n 5), and 24 threads following the recommendations in the user guide.

### 2.11 Identification of Differentially Expressed Genes (DEGs)

Transcript quantification was performed with Salmon (Patro *et al*., 2017). After indexing the *de novo* transcriptome, quantification was run with --*validateMappings* and *--gcBias* options using 24 threads. Raw counts were imported into RStudio using *tximport* (Soneson *et al*., 2015) and analyzed within a *DGEList* object in edgeR (Robinson *et al*., 2010). Lowly expressed genes were filtered using *filterByExpr* (Chen *et al*., 2016), and counts were normalized by the Trimmed Mean of M-values (TMM) method. Differential expression analysis was conducted using edgeR’s negative binomial models, with p-values adjusted by the Benjamini-Hochberg method; clusters with FDR < 0.05 were considered significantly differentially expressed. Functional annotation of genes was performed using BLAST (Camacho *et al*., 2009) against the NCBI nr (non-redundant protein) and nt (nucleotide) databases with an e-value < 1×10^−6^. BLASTX searches against nr were conducted for differentially expressed genes (DEGs) using BioData Hub (Department of Biology, University of Padova, Italy) resources and used as the primary source for downstream annotation. Hits were filtered by collapsing redundant matches, assigning taxonomic information with taxoniq, and removing queries whose top three nr/nt hits were not assigned to Streptophyta. For each gene, the top five nr hits were retained based on e-value and bitscore. Final annotations were assigned by prioritizing database sources in the following order: RefSeq → UniProt ID (PDB/Swiss-Prot/TrEMBL) → others.

UniProt identifiers were retrieved through the UniProt ID mapping service. To assign a single representative entry per cluster, duplicates were first collapsed by query-UniProt pairs and subsequently by protein name, retaining entries associated with the highest number of Gene Ontology (GO) terms; unresolved cases were manually curated. When the best nr hit lacked a UniProt ID, alternative hits among the top five were manually evaluated, and the most appropriate annotation was selected.

### 2.12 Gene Ontology analysis

Gene Ontology (GO) enrichment analysis was performed in R (v4.5.1) using the package topGO (Alexa & Rahnenführer, 2025), with UniProt identifiers as proxies for genes (FDR <0.01). Given the absence of a reference genome, the background gene set was defined using the UniProt proteome of *Eragrostis curvula* (Poaceae), a C_4_ species phylogenetically close to *Sporobolus* (Huang *et al*., 2022), and well annotated in public databases (*e*.*g*., QuickGO). Enrichment was tested using the *classic* algorithm with Fisher’s exact test, and p-values were adjusted for multiple testing using the Benjamini-Hochberg procedure. Only GO terms at least three levels below the root within the Biological Processes (BP), Cellular Components (CC), and Molecular Functions (MF) categories were considered. Terms with a FDR < 0.05 were deemed significantly enriched.

### 2.13 Untargeted metabolomics

Leaf samples (20 mg Fresh Weight/FW; three biological replicates per species) were homogenized in 400 µL HPLC-grade methanol using a TissueLyser LT (Qiagen, Germany; 50 Hz, 15 s), centrifuged (5’000 × g, 5 min), and supernatants analyzed by UHPLC-HRMS (1290 Infinity II LC system coupled to a G6550B Q-TOF mass spectrometer; Agilent Technologies, USA) with Dual AJS ESI source (Agilent Technologies, USA) in positive ion mode. Extracts (5 µL) were separated on a ZORBAX SB-C18 RRHT column (3.0 × 50 mm, 1.8 µm) (Agilent Technologies, USA) using water (phase A) and acetonitrile (phase B), both with 0.1% formic acid, at 0.6 mL/min. Chromatographic separation was carried out using a linear gradient of solvent B in solvent A, as follows: the initial mobile phase composition (90% A and 10% B, v/v) was maintained for 3 min, then solvent B was linearly increased to 45% over 20 min and further to 90% at 30 min, then re-equilibration to initial conditions. MS data were acquired in AutoMS² mode (m/z 100-3000) with MassHunter software (Agilent Technologies, USA).

Source parameters were: gas temperature 210 °C; gas flow 20 L/min; nebulizer 35 psi; sheath gas temperature 250 °C; sheath gas flow 12 L/min; capillary voltage (VCap) 3500 V; fragmentor 250 V; nozzle voltage 1000 V. LC-MS output files were processed according to Trentin *et al*. (2025). Processed data were analyzed: (1) in Global Natural Product Social Molecular Networking (GNPS) for library matching and feature-based molecular networking (FBMN) (Table S3 and S4) (Nothias *et al*., 2020); (2) in SIRIUS according to Trentin *et al*. (2025). GNPS and SIRIUS annotations were merged, manually curated using literature data, and confidence levels were assigned following Sumner *et al*. (2007).

Statistical analyses were performed in R using *ggsci*, *matrixStats*, *ggrepel*, and *tidyverse*. Significance was tested by one-way ANOVA followed by Tukey’s post-hoc test. Features were visualized with heatmaps using *pheatmap* (row mean normalization; Euclidean clustering). Variables contributing most to group separation were identified with Random Forest in MetaboAnalyst (Pang *et al*., 2024) using variable importance scores (VIP).

### 2.14 Statistical analyses

Except for Gene Ontology enrichment and metabolomics analyses (see above), all statistical analyses were performed in R (v4.5.1) using one-way ANOVA followed by Tukey’s post-hoc test. All data were obtained from at least three independent biological replicates and are presented as mean ± standard deviation (SD). Statistical significance was set at p-value < 0.05. Details of statistical tests and significance levels are provided in the corresponding figure captions.

## 3. Results

### 3.1 Survival analysis

The native species’ (*S. maritimus*) survival rate decreased after the end of the heatwave, with the lowest value (25%) recorded at the end of the experiment (Figure 2). In contrast, the NIS (*S. anglicus*) showed no change in survival rate, with only one plant dying throughout the entire experimental period. Apparently, in terms of survival, the native species might not tolerate this type of stress as well as the NIS.

**Figure 2.**
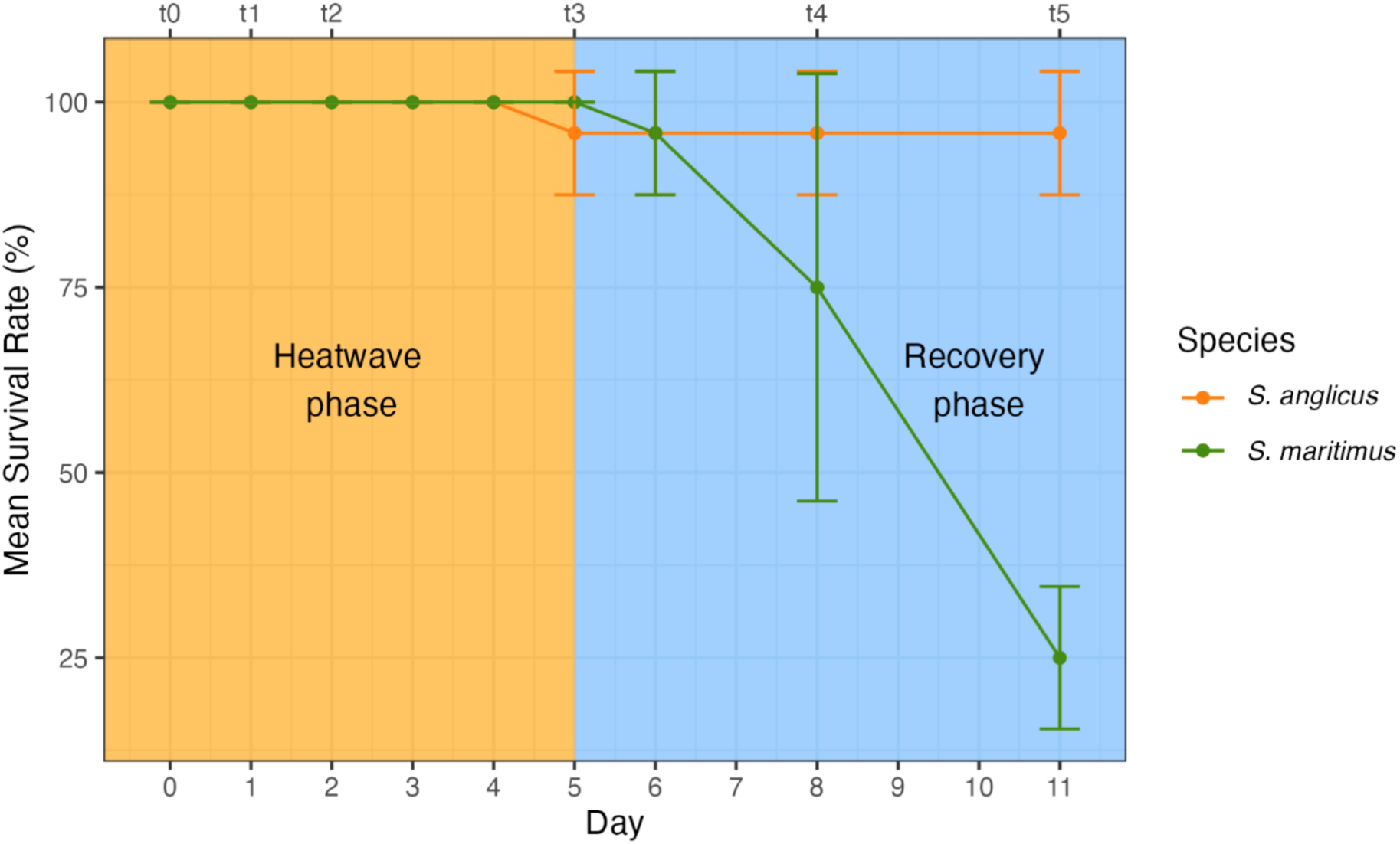
Survival rates during heatwave and recovery phases. The graph shows mean survival (± SD), expressed as the percentage of living plants for *S. maritimus* (green line) and *S. anglicus* (orange line). The orange box indicates the heatwave phase (t_0_-t_3_), the light blue box the recovery phase (t_3_-t_5_).

### 3.2 Ultrastructural changes

Alteration of organization of the thylakoid membranes was examined before, during, and after heatwave exposure using TEM (Figure 3).

**Figure 3.**
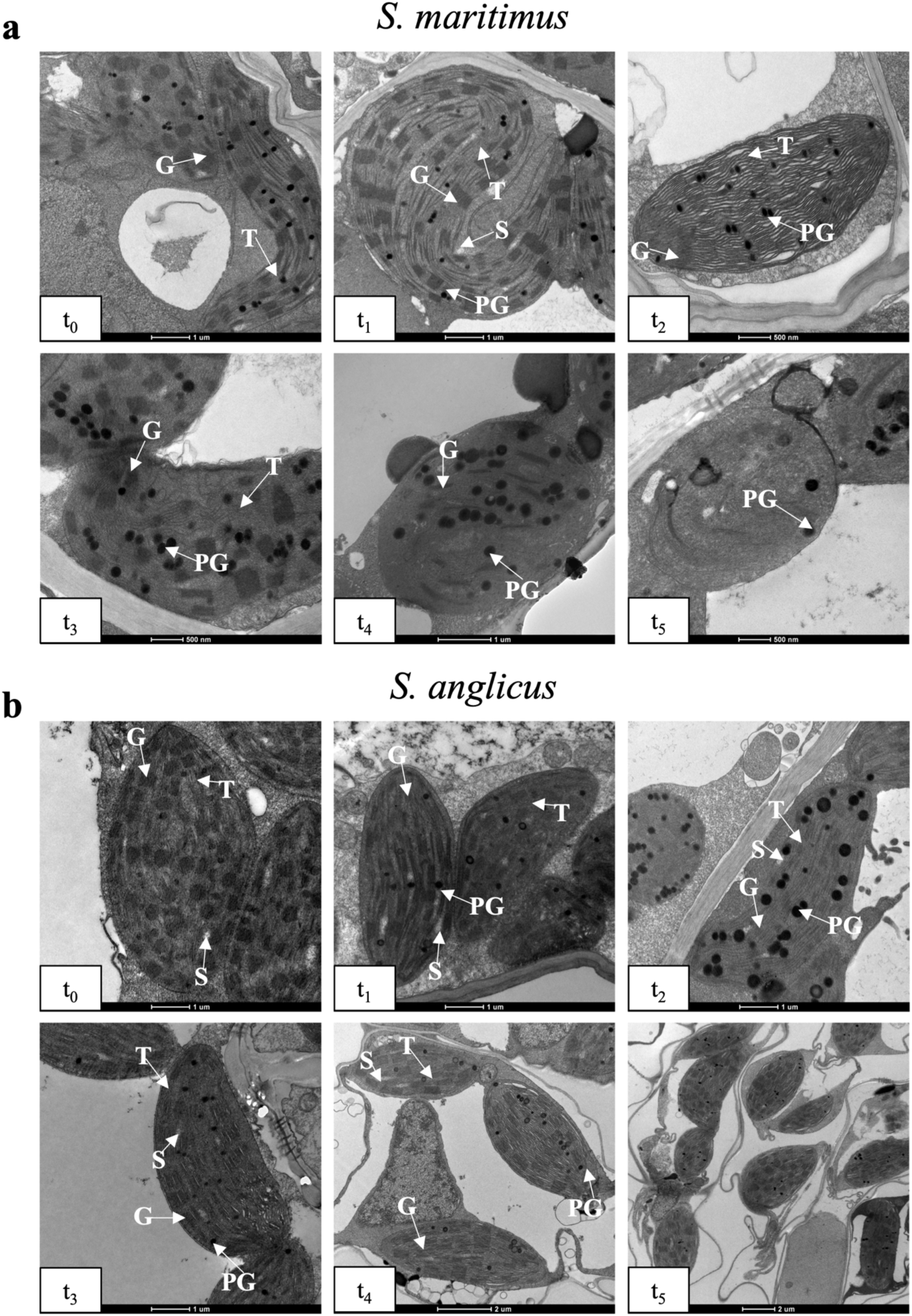
Ultrastructure of the chloroplast. Chloroplast ultrastructure of *S. maritimus* (**a**) and *S. anglicus* (**b**) before, during, and after heat stress. G = grana, T = thylakoid membranes, S = starch, PG = plastoglobuli. Scale bars are shown below each image for reference.

Changes in chloroplast organization were evident in both species as early as t_2_. Clear differences between the species emerged in *S. maritimus* (native), where evident dilation of thylakoid membranes (T) and scarce grana (G) were present (Figure 3a); in *S. anglicus* (NIS), instead, thylakoids remained stacked forming evident grana, but with a notable increase in plastoglobuli (PG) (Figure 3b). The differences were more pronounced with increasing time. At t_3_, the native species exhibited unstacking of thylakoids and some grana, but above all a high accumulation of PG. In contrast, the NIS showed only some dilatations of the thylakoids and not many grana. During the recovery phase, damage in the native species further progressed, leading to the degeneration of internal chloroplast organization. Conversely, the NIS showed signs of recovery as early as t_4_, and, by t_5_, chloroplast ultrastructure closely resembled the initial state (t_0_).

In summary, the native species showed more pronounced alterations after just 36 hours of exposure, while the NIS demonstrated better stress tolerance even after 5 days of exposure. Furthermore, the native species showed no recovery in chloroplast ultrastructure as temperatures declined, whereas *S. anglicus* exhibited clear signs of it. These findings suggest that *S. anglicus* has greater tolerance to heat stress and higher recovery capacity compared to *S. maritimus*, at least with respect to chloroplast structure.

### 3.3 Photosynthetic apparatus efficiency and pigment content variation

Disorganization of photosynthetic membranes is often associated with reduced photosynthetic efficiency, reflecting a lower capacity to convert light energy into chemical energy.

Considering F_v_’/F_m_’ (Figures 4a and b), both species showed a decline from t_0_ to t_3_. At t_3_, *S. maritimus* (Figure 4a) reached lower values compared to *S. anglicus* (Figure 4b). Additionally, the native species’ values continued to decrease after t_3_, with only a slight recovery at t_5_ (Figure 4a). In contrast, *S. anglicus* (Figure 4b) displayed an increase in F_v_’/F_m_’ at t_4_ and t_5_. The same pattern was observed for the actual efficiency of PSII (ϕII) (Figures 4c and d). At the same time, ϕNPQ increased during the stress phase in both species up to t_3_, with a more pronounced rise in *S. maritimus* (Figure 4e). After the end of the heatwave (t_3_), *S. maritimus* showed a further increase at t_4_ followed by a slight decrease at t_5_ (Figure 4e), whereas in *S. anglicus ϕ*NPQ decreased from t_4_ onward and continued to decline until t_5_ (Figure 4f).

**Figure 4.**
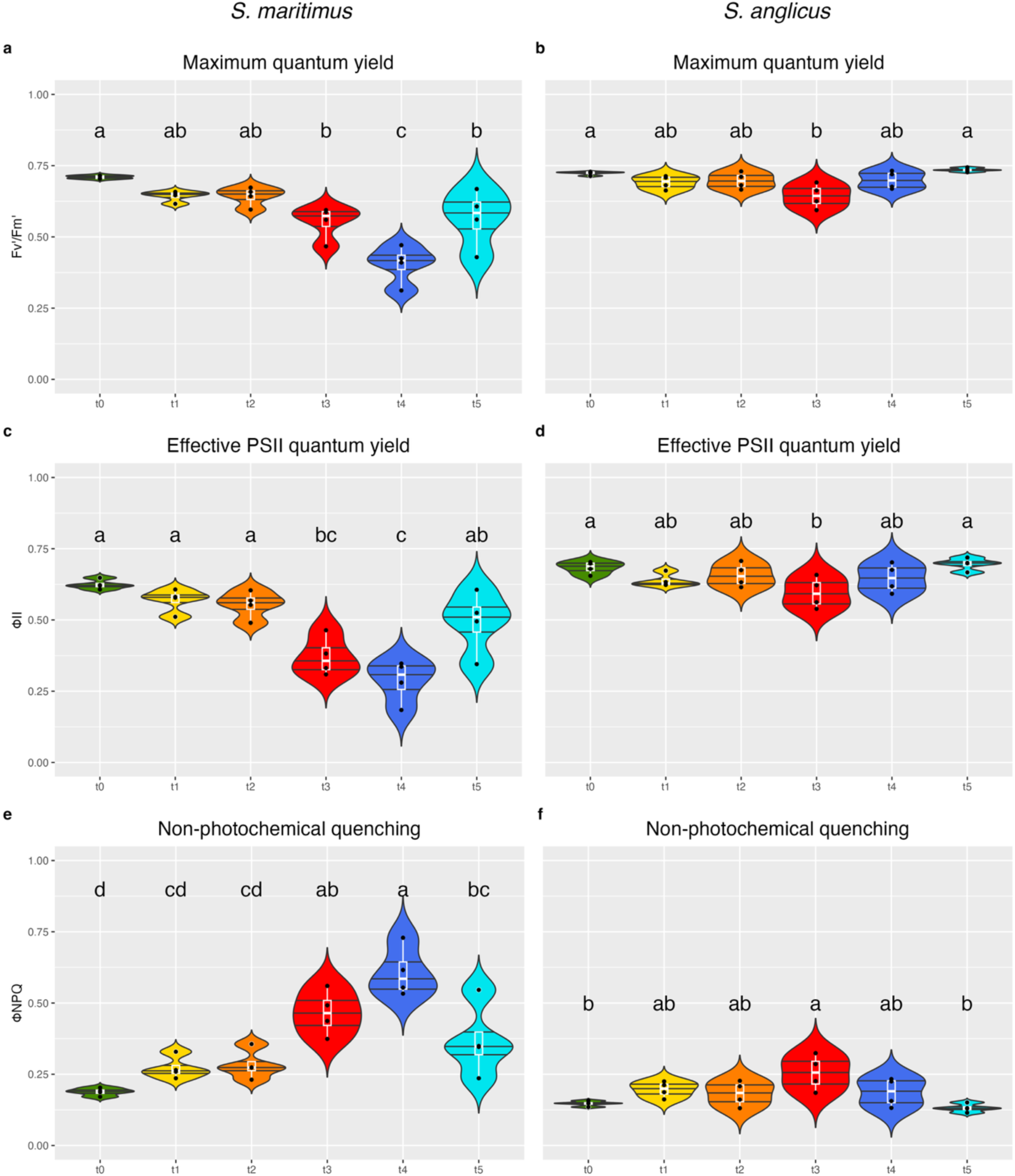
Photosynthetic efficiency variation before, during, and after the heatwave stress. Variation in maximum quantum yield (F_v_’/F_m_’) (**a**-**b**), effective PSII quantum yield (ϕPSII) (**c**-**d**), and non-photochemical quenching (ϕNPQ) (**e-f**) in *S. maritimus* (**a**, **c**, **e**) and *S. anglicus* (**b**, **d**, **f**) from t_0_ to t_5_. The results are expressed as mean ± SD (n≥3). One-way ANOVA with Tukey’s post-hoc test was carried out to determine significant differences; results are reported in Tables S5 and S6. For each plot, means with different letters showed significant differences among different time points.

In summary, both species experienced a decline in photosynthetic efficiency (F_v_’/F_m_’ and ϕII) under increasing stress, accompanied by a corresponding rise in energy dissipation (ϕNPQ). However, striking differences emerged during recovery: the native species (*S. maritimus*) failed to restore photosynthetic parameters to their initial levels, whereas the NIS (*S. anglicus*) fully recovered, highlighting its superior resilience to heat stress.

Since photosynthetic efficiency is strictly correlated with pigment content, chl *a*, chl *b*, and carotenoids were measured during the different phases of the experiment.

The native species showed a significant decrease of total chlorophylls content starting from t_4_, with the lowest value at t_5_ (Figure 5a), whilst the NIS did not show any significant variation (Figure 5b). In particular, chl *a* and chl *b* decreased from t_4_ in the native species (Figures 5c and 5e). In the NIS, chl *a* did not vary and chl *b* was significantly different only in the comparison t_1_ *vs.* t_4_ (Figures 5d and 5f). Instead, the ratio between chl *a* and chl *b*, which is linked to LHCs size (Tanaka & Tanaka, 2011), decreased significantly at t_5_ compared to t_0_ in both species (Figures 5g and 5h).

**Figure 5.**
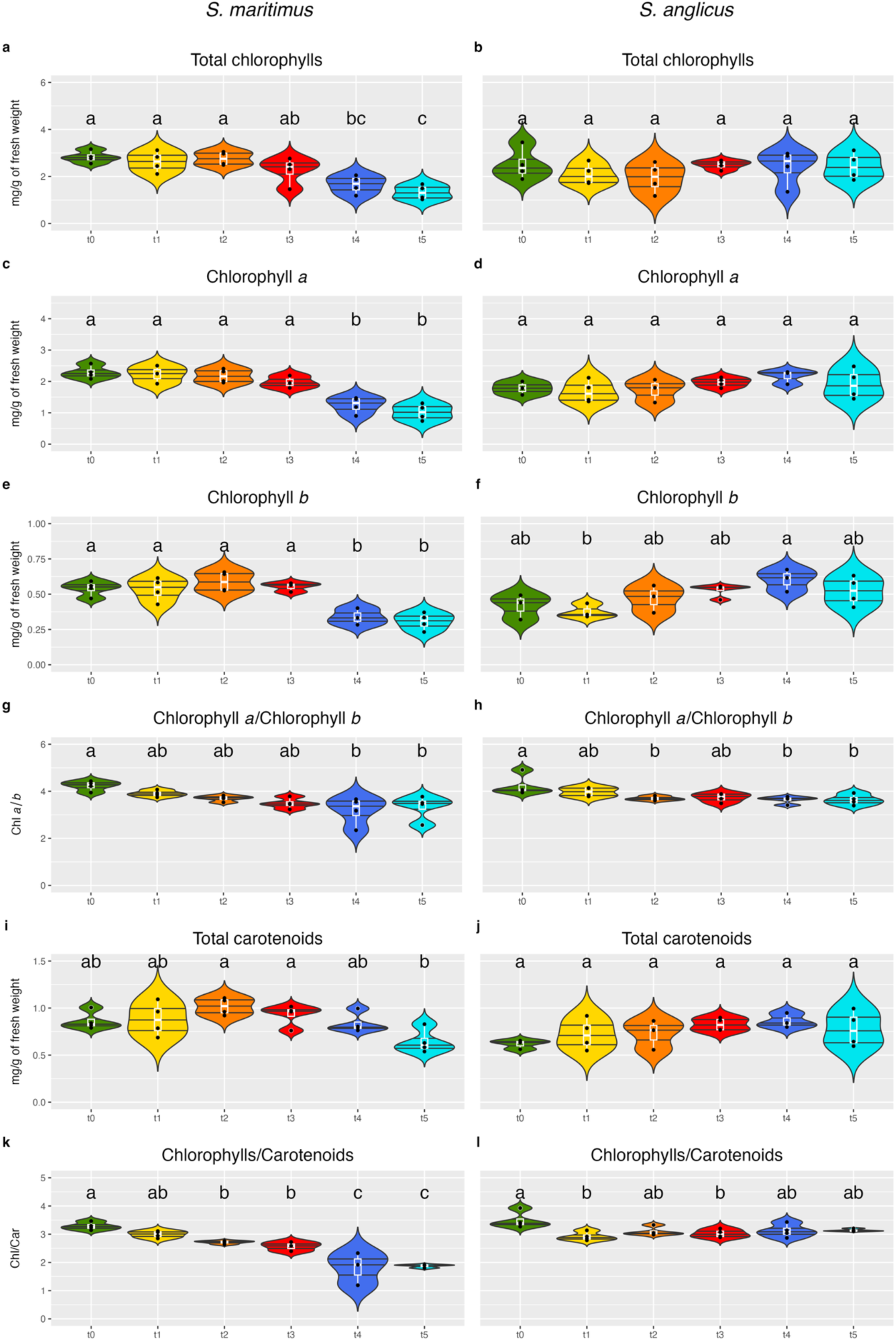
Pigment content variation before, during, and after the heatwave stress. Variation in Total Chlorophylls (**a-b**), Chlorophyll *a* (**c-d**), Chlorophyll *b* (**e-f**), Chlorophyll *a*/Chlorophyll *b* (**g-h**), Total Carotenoids (**i-j**), and Chlorophylls/Carotenoids (**k-l**) in *S. maritimus* (**a, c, e, g, i, k**) and *S. anglicus* (**b, d, f, h, j, l**) from t_0_ to t_5_. The results (n≥3) are expressed as mean ± SD. One-way ANOVA with Tukey’s post-hoc test was carried out to determine significant differences; results are reported in Tables S5 and S6. For each plot, means with different letters showed significant differences among different time points.

Total carotenoids, which compose LHCs and can function as antioxidants (Wang *et al*., 2018), decreased significantly at t_5_ compared to t_2_ and t_3_ in *S. maritimus* (Figure 5i), with no variation in *S. anglicus* (Figure 5j). The profiles of the most abundant carotenoids, measured by HPLC, are presented in Figure S3. The decrease of the chl *a*+*b*/carotenoids (chl/car) ratio can be used as a stress index (Esteban *et al*., 2015). It decreased significantly in *S. maritimus* from t_0_ to t_5_ (Figure 5k). On the other hand, in *S. anglicus* we observed significant differences with t_0_ only at t_1_ and t_3_ (Figure 5l).

Overall, the native species experienced greater damage to photosynthetic efficiency and pigment content compared to the NIS, as exemplified by the pronounced decreases in F_v_’/F_m_’, chlorophylls, and carotenoids, as well as the lack of recovery in both photosynthetic efficiency parameters and the chl/car ratio, which instead were observed in *S. anglicus*.

### 3.4 Antioxidant enzyme activity changes

In order to evaluate plant responses to oxidative stress, the activities of key ROS-scavenging enzymes were detected. Ascorbate peroxidase (APx) activity increased significantly in *S. maritimus* at t_4_ and remained elevated at t_5_ (Figure 6a), whereas in *S. anglicus* it peaked at t_3_ and returned to t_0_ levels at t_4_ and t_5_ (Figure 6b). Guaiacol peroxidase (GPx) activity in *S. maritimus* increased significantly only at t_5_ (Figure 6c), while in *S. anglicus* the highest activity was observed at t_2_, followed by a return to baseline values (Figure 6d). Catalase (CAT) activity in native species increased steadily after t_2_, with significantly higher values at t_5_ compared to earlier time points (Figure 6e); no significant variation was detected in the NIS (Figure 6f). Superoxide dismutase (SOD) activity did not vary significantly in either species throughout the experiment (Figures 6g and 6h). Copper chelating activity (CCA) remained unchanged until t_3_ in both species, increasing significantly only at t_5_ in *S. maritimus*, whereas in *S. anglicus* it increased at t_3_ and returned to baseline at t_5_ (Figures 6i and 6j).

**Figure 6.**
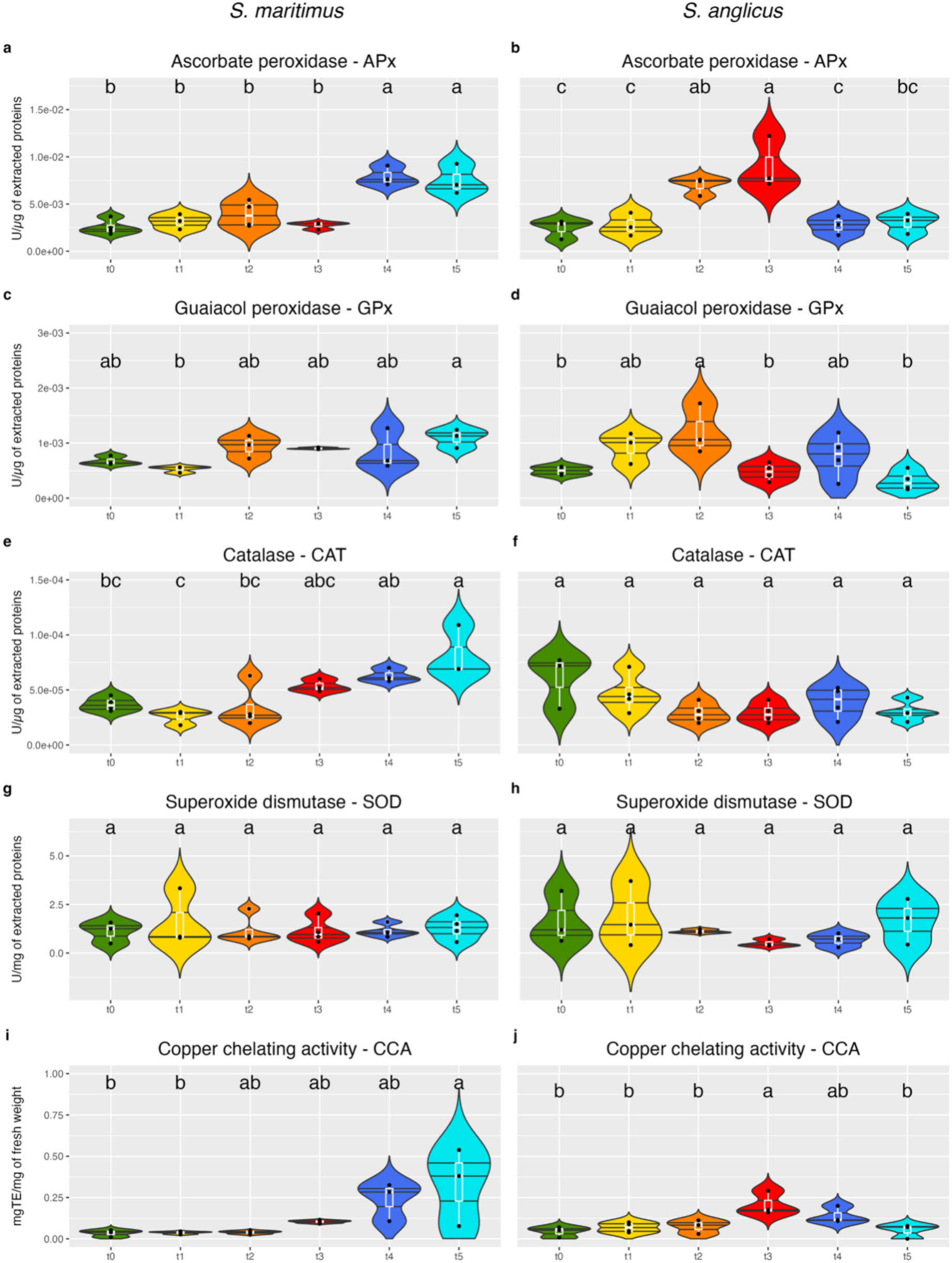
Antioxidant parameters variation before, during, and after the heatwave stress. Variation in Ascorbate Peroxidase (APx) (**a-b**), Guaiacol Peroxidase (GPx) (**c-d**), Catalase (CAT) (**e-f**), Superoxide dismutase (SOD) (**g-h**), and Copper Chelating Activity (CCA) (**i-j**) in *S. maritimus* (**a, c, e, g, i**) and *S. anglicus* (**b, d, f, h, j**) from t_0_ to t_5_. The results (n≥3) are expressed as mean ± SD. One-way ANOVA with Tukey’s post-hoc test was carried out to determine significant differences; results are reported in Tables S5 and S6. For each plot, means with different letters showed significant differences among different time points.

In conclusion, antioxidant defenses in the native species responded slowly, peaking during the recovery phase rather than during the heatwave. In contrast, the NIS showed a progressive increase in both antioxidant activity and metal-chelating capacity as stress intensified.

### 3.5 Gene expression analyses

To better clarify how the two plant species respond to heat stress and to uncover the mechanisms underlying their differences, we generated a *de novo* transcriptome for the genus *Sporobolus* and performed gene expression analyses on samples collected throughout the simulated heatwave experiment.

#### 3.5.1 *Sporobolus de novo* transcriptome assembly

*De novo* transcriptome assembly yielded 89’523 transcripts for *S. maritimus* and 113’311 for *S. anglicus*. After removal of non-coding RNA fragments and redundancy reduction (see Materials and Methods), 42’947 and 52’180 non-redundant transcript clusters were retained, respectively. The longest sequence per cluster was used for downstream analyses. Assembly quality was supported by N50 values of 1’111 bp for *S. maritimus* and 1’147 bp for *S. anglicus* and by high read mapping rates (92.3% and 89.4%, respectively).

#### 3.5.2 Identification and annotation of DEGs after the simulated heatwave

To assess transcriptional responses to heat stress, RNA-seq analysis was performed on 36 biological replicates (3 per time point per species), excluding three samples (two *S. maritimus* and one *S. anglicus*) due to technical issues. Principal component analysis (PCA) revealed clear clustering of samples by experimental phase (Figure 7). In both species, samples collected during the heatwave phase grouped together. In *S. maritimus*, early time points (t_1_ and t_2_) were clearly separated from subsequent stages, with t_3_ showing a distinct expression profile (Figure 7a). In *S. anglicus*, samples from t_1_ to t_3_ clustered more closely than in *S. maritimus*, although they remained distinct from later time points (Figure 7b).

**Figure 7.**
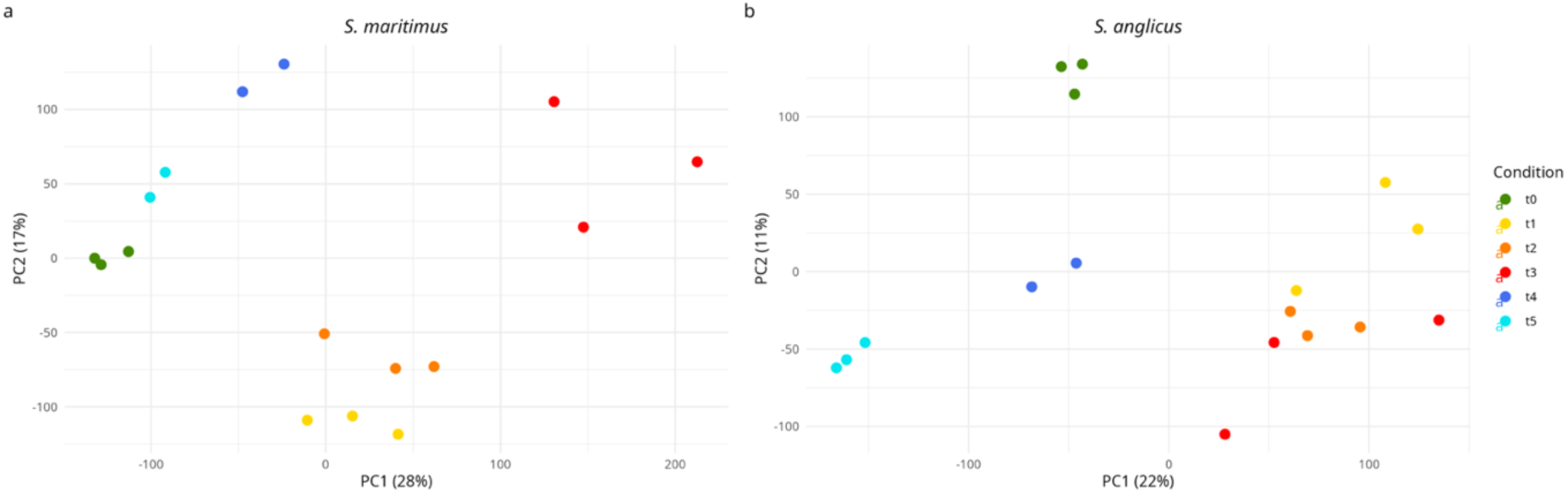
Principal Component Analysis (PCA) of gene expression profiles during the simulated heatwave experiment. All biological replicates used for DEGs detection are shown for *S. maritimus* (**a**) and *S. anglicus* (**b**). Different colours indicate different time points: (1) green - t_0_; (2) yellow - t_1_; (3) orange - t_2_; (4) red - t_3_; (5) blue - t_4_; (6) light blue - t_5_.

Based on the PCA results, differentially expressed genes were identified (t_0_ *vs*. t_1_, t_0_ *vs*. t_3_, t_3_ *vs*. t_5_ and t_0_ *vs.* t_5_) and subsequently annotated. The native species exhibited a higher number of DEGs (FDR < 0.01) than the non-indigenous species in all comparisons except t_0_ *vs*. t_5_ (Figure S4). Particularly, the number of DEGs in the native species was 3’470 (8.08%) in t_0_ *vs*. t_1_, 11’049 (25.73%) in t_0_ *vs*. t_3_, 7’744 (18.03%) in t_3_ *vs*. t_5_ and 171 (0.40%) in t_0_ *vs.* t_5_. In the NIS we observed 1’816 (3.48%) in t_0_ *vs*. t_1_, 2’034 (3.90%) in t_0_ *vs*. t_3_, 6’005 (11.50%) in t_3_ *vs*. t_5_ and 1’594 (3.05%) in t_0_ *vs.* t_5_. Across all comparisons, up- and down-regulated genes were observed in similar proportions. Full DEG lists are provided in Tables S7-S14.

Functional annotation assigned UniProt identifiers to 15’979 unique clusters in *S. maritimus* (∼75% of all identified DEGs) and 12’953 in *S. anglicus* (∼78%) following automated annotation and manual curation (see Materials and Methods). These annotated clusters were used for downstream Gene Ontology (GO) enrichment analyses.

#### 3.5.3 Gene Ontology (GO) analysis

GO enrichment analysis was performed to identify Biological Processes (BP), Molecular Functions (MF), and Cellular Components (CC) categories associated with differential expression in *S. maritimus* and *S. anglicus*. Analyses were conducted for the short-term heat stress (t_0_ *vs.* t_1_, Figure 8), the long-term heat stress (t_0_ *vs.* t_3_, Figure 9), the recovery phase (t_3_ *vs.* t_5_, Figure 10) and the comparison with initial conditions (t_0_ *vs.* t_5_, Figure S5). Full lists of enriched GO terms for each species and comparison are provided in Tables S15-21.

**Figure 8.**
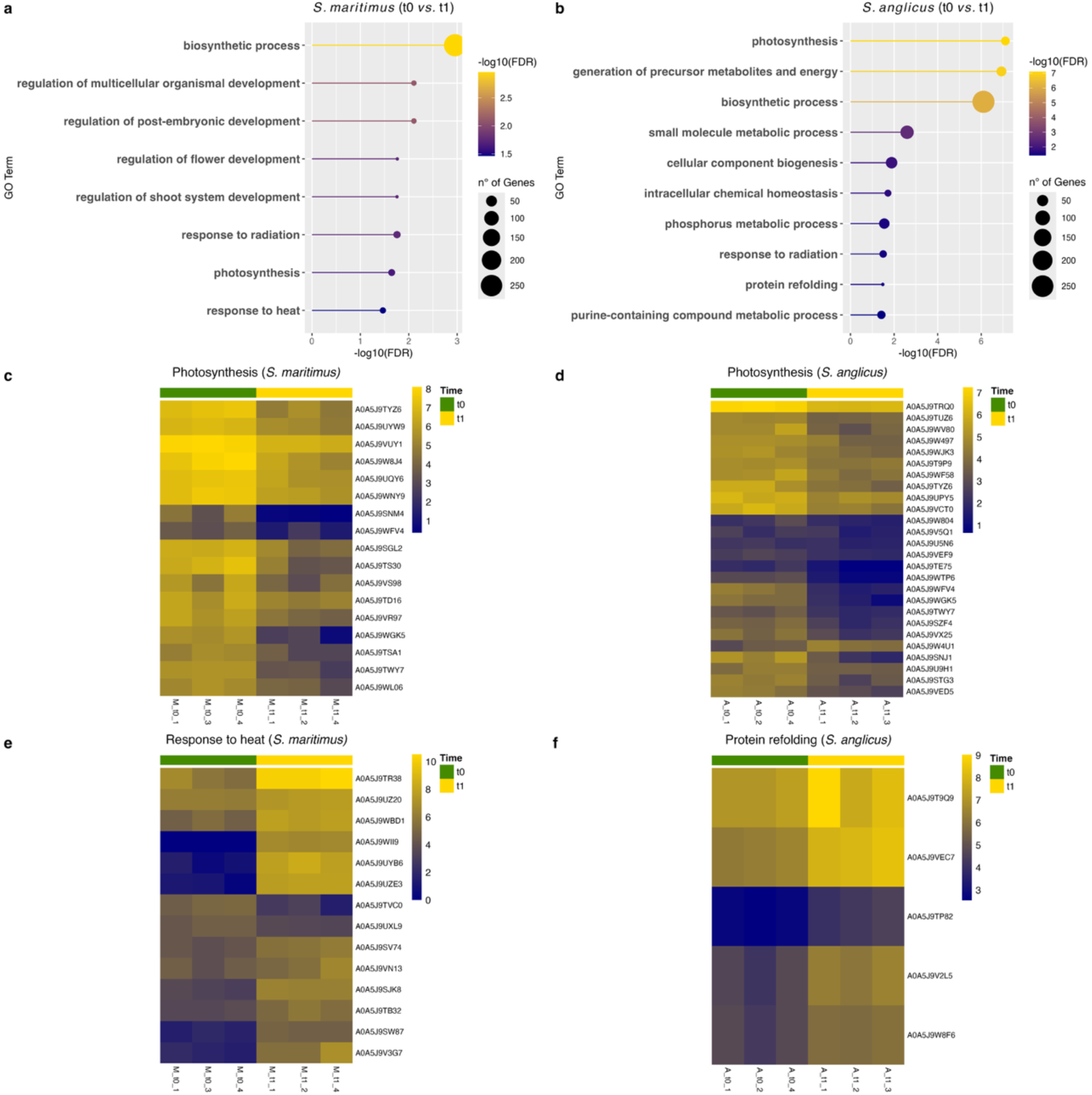
GO analysis of differentially expressed genes after short-term heat stress (t_0_ *vs*. t_1_). (**a–b**) Most enriched Biological Process (BP) terms in *S. maritimus* and *S. anglicus*, respectively; circle size is proportional to the number of DEGs and color (blue-low, yellow-high) is proportional to the statistical significance, expressed as the negative logarithm of the False Discovery Rate (-log10(FDR)). (**c–f**) Heatmaps of expression values for genes contributing to specific BP terms: “Photosynthesis” (GO:0015979) for *S. maritimus* (**c**) and *S. anglicus* (**d**); “Response to heat” (GO:0009408) for *S. maritimus* (**e**); “Protein refoldin*g*” (GO:0042026) for *S. anglicus* (**f**). The UniProt identifier is reported on each row. The color of the cell is proportional to the expression value (blue-low, yellow-high), represented as log2. Complete DEGs lists and enriched GO terms for both species are provided in Tables S7 and S15 (*S. maritimus*) and S11 and S18 (*S. anglicus*).

**Figure 9.**
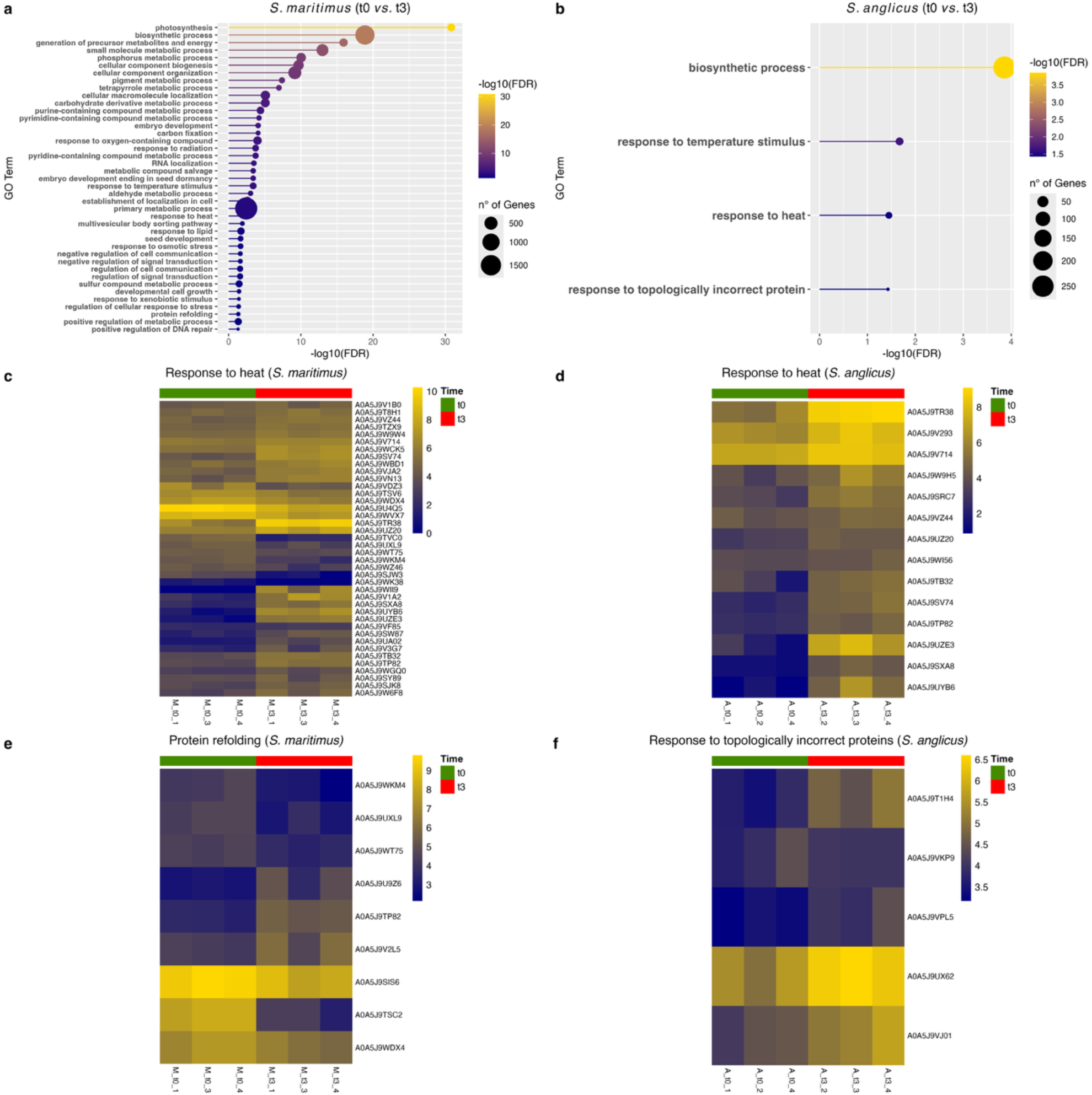
GO analysis of differentially expressed genes after long-term heat stress (t_0_ *vs*. t_3_). (**a–b**) Most enriched Biological Process (BP) terms in *S. maritimus* and *S. anglicus*, respectively; circle size is proportional to the number of DEGs and color (blue-low, yellow-high) is proportional to the statistical significance, expressed as the negative logarithm of the False Discovery Rate (-log10(FDR)). (**c–f**) Heatmaps of expression values for genes contributing to specific BP terms: “Response to heat” (GO:0009408) for *S. maritimus* (**c**) and for *S: anglicus* (**d**); “Protein refolding” (GO:0042026) for *S. maritimus* (**e**); “Response to topologically incorrect proteins” (GO:0035966) for *S. anglicus* (**f**). The UniProt identifier is reported on each row. The color of the cell is proportional to the expression value (blue-low, yellow-high), represented as log2. Complete DEGs lists and enriched GO terms for both species are provided in Tables S8 and S16 (*S. maritimus*) and S12 and S19 (*S. anglicus*).

**Figure 10.**
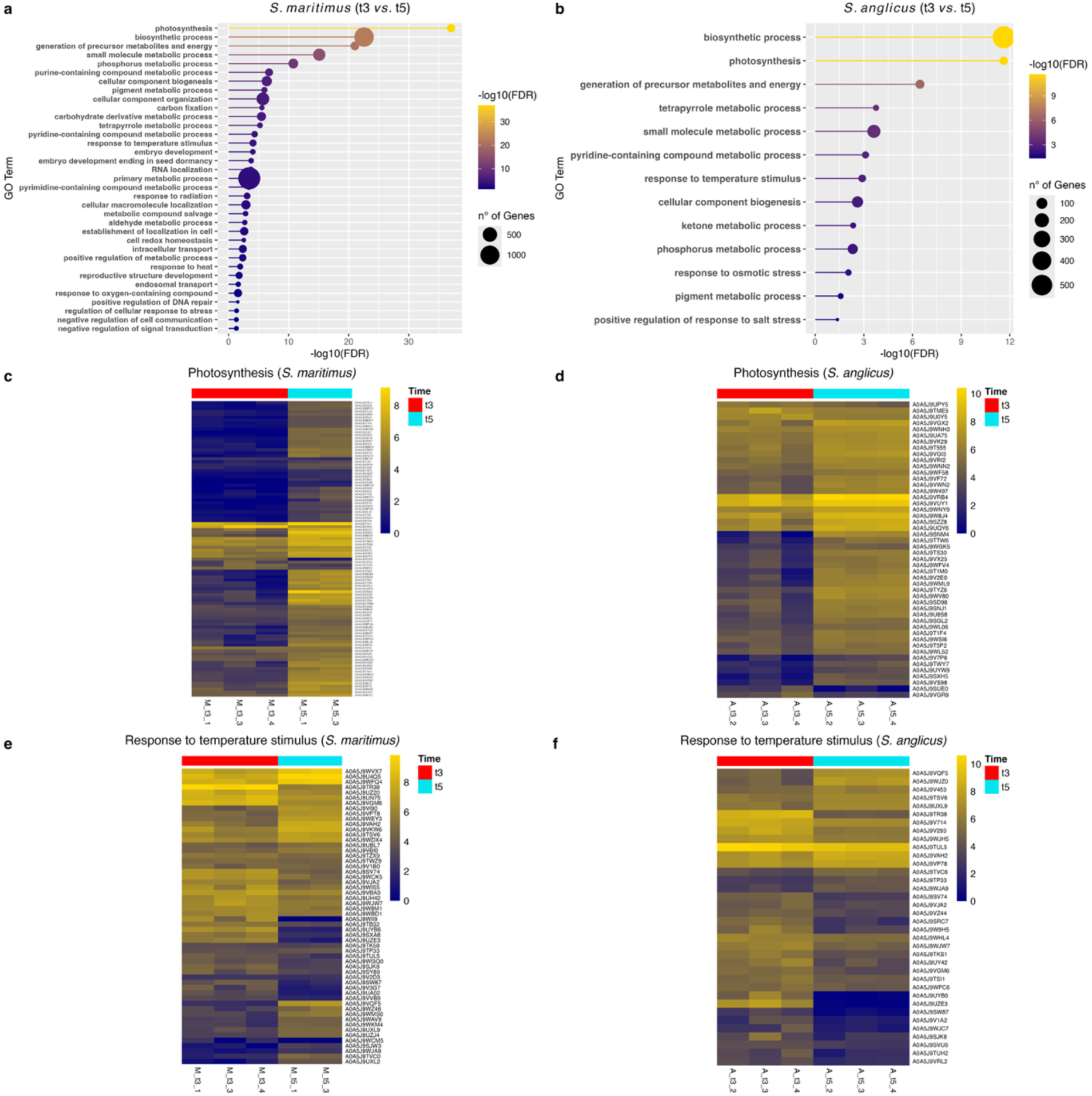
GO analysis of differentially expressed genes after recovery phase (t_3_ *vs*. t_5_). (**a–b**) Most enriched Biological Process (BP) terms in *S. maritimus* and *S. anglicus*, respectively; circle size is proportional to the number of DEGs and color (blue-low, yellow-high) is proportional to the statistical significance, expressed as the negative logarithm of the False Discovery Rate (-log10(FDR)). (**c–f**) Heatmaps of expression values for genes contributing to specific BP terms: “Photosynthesis” (GO:0015979) for *S. maritimus* (**c**) and *S. anglicus* (**d**); “Response to temperature stimulus” (GO:0009266) for *S. maritimus* (**e**) and for *S. anglicus* (**f**). The UniProt identifier is reported on each row. The color of the cell is proportional to the expression value (blue-low, yellow-high), represented as log2. Complete DEGs lists and enriched GO terms for both species are provided in Tables S9 and S17 (*S. maritimus*) and S13 and S20 (*S. anglicus*).

##### Short-term heat stress response

During short-term stress (t_0_ *vs.* t_1_, Figure 8), the BP term “photosynthesis” was significantly enriched in both species (Figures 8a and 8b), with nearly all associated genes being downregulated (17/17 in *S. maritimus*, 25/26 in *S. anglicus*, Figures 8c and 8d). However, only the native species (Figure 8a) showed enrichment of thermal stress-related BP terms, such as “response to heat” (Figure 8e), along with effects on developmental processes (*e.g.*, “regulation of multicellular organism development”). In contrast, in the NIS (Figure 8b), upregulated DEGs were enriched for “protein refolding” (Figure 8f). At the MF level (Figure S6), *S. maritimus* showed enrichment of protein-binding terms associated with chaperonins, including “protein-folding chaperone binding” (9/9 genes upregulated) and “heat shock protein binding” (8/9 genes upregulated). Both species displayed enrichment of the “unfolded protein binding” term; however, in the native species most genes were upregulated (10/12), whereas in the NIS all associated genes were upregulated (12/12) (Figures S7a and b).

##### Long-term heat stress response

After 5 days of heat stress (t_0_ *vs*. t_3_, Figure 9), *S. maritimus* showed an increased number of enriched BP terms, accompanied by a higher number of DEGs, indicating a broader impact on multiple biological processes. Photosynthesis-related processes, including “photosynthesis”, “pigment metabolic process”, and “carbon fixation” were predominantly downregulated (Figures 9a, S8a, and S8b). Although “response to heat” and “response to temperature stimulus” remained significantly enriched (Figure 9a), a subset of the associated genes was downregulated (12/40 and 25/65, respectively; Figure 9c). Notably, “protein refolding” was also enriched at this time point in the native species (Figure 9e). Terms related to ROS, such as “response to oxygen-containing compounds” were also enriched (Figure 9a), including three L-ascorbate peroxidase genes, two of which were upregulated. In the NIS, the number of DEGs at t_0_ *vs.* t_3_ (2’034 DEGs) was comparable to that observed at t_0_ *vs.* t_1_ (1’816 DEGs). Enriched BP terms included “response to temperature stimulus”, “response to heat”, and “response to topologically incorrect proteins”, with all associated genes upregulated (Figures 9b, 9d, and 9f). For MF terms (Figure S6), protein-binding functions were enriched in both species. In *S. maritimus* (Figure S7c), genes contributing to “unfolded protein binding” were partially downregulated (13/34), whereas in *S. anglicus* nearly all genes were upregulated (Figure S7d). Considering the CC category (Figures S9c and S9d), *S. maritimus* showed enrichment of protein degradation-related terms (“proteasome complex”, “peptidase complex”), with most genes upregulated (27/29 and 34/40, respectively), alongside photosynthetic apparatus terms (“photosystem”), with most genes being downregulated (42/44).

##### Recovery effect after long-term heat stress

Six days after the end of heat stress (t_3_ *vs.* t_5_, Figure 10), both species (Figures 10a and 10b) exhibited upregulation of genes associated with the BP terms “photosynthesis” (Figures 10c and 10d) and “pigment metabolic processes”. The BP term “response to temperature stimulus” was also enriched in both species, but most genes were downregulated (31/56 in *S. maritimus* and 26/36 in *S. anglicus*, Figures 10e and 10f). In *S. maritimus*, additional enrichment was observed for “response to oxygen-containing compounds” and “regulation of cellular response to stress”, with approximately half of the genes still upregulated. In *S. anglicus*, recovery of the CC term related to Oxygen Evolving Complex (OEC) was evident, with 8 out of 10 genes in the “Photosystem II oxygen-evolving complex” term being upregulated (Figure S9f).

Finally, to assess the lasting effects of the simulated heatwave, we examined the t_0_ *vs.* t_5_ comparison (Figures S5a, b and c). Due to the high mortality observed in the native species, this analysis was performed only for *S. anglicus*. Enriched BP terms included those related to cell development (*e.g.*, “cellular component biogenesis”) and photosynthesis (“photosynthesis” and “pigment metabolic process”) (Figure S5a). Most associated genes were upregulated (31/47 for “cellular component biogenesis”, 23/26 for “photosynthesis”, and 15/16 for “pigment metabolic process”).

### 3.6 Metabolites variation

A total of 506 features were detected from *S. maritimus* and *S. anglicus*. For each feature, mass-to-charge ratio, retention time, adduct type, putative molecular formula, annotation level, annotated name (for level 2 identifications), and compound classification are reported in Table S22. During heat stress, a greater number of metabolite features varied in *S. maritimus* (61) than in *S. anglicus* (31). Despite the high proportion of unannotated features (annotation level 4), analysis of the 50 most significant features revealed a pronounced shift in metabolite profiles in *S. maritimus* after 5 days of exposure (t_3_, Figure 11a), whereas changes were less marked in *S. anglicus* (Figure 11b). Partial Least Squares Discriminant Analysis (PLS-DA) with Variable Importance in Projection (VIP) scatter plots highlighted the features contributing most to class separation (Figure 11c-d). In *S. maritimus*, these features increased progressively with stress, a pattern not observed in the NIS. Overall, despite the large proportion of unannotated features, the native species exhibited a more pronounced shift in metabolite composition in response to heat stress compared to the NIS.

**Figure 11.**
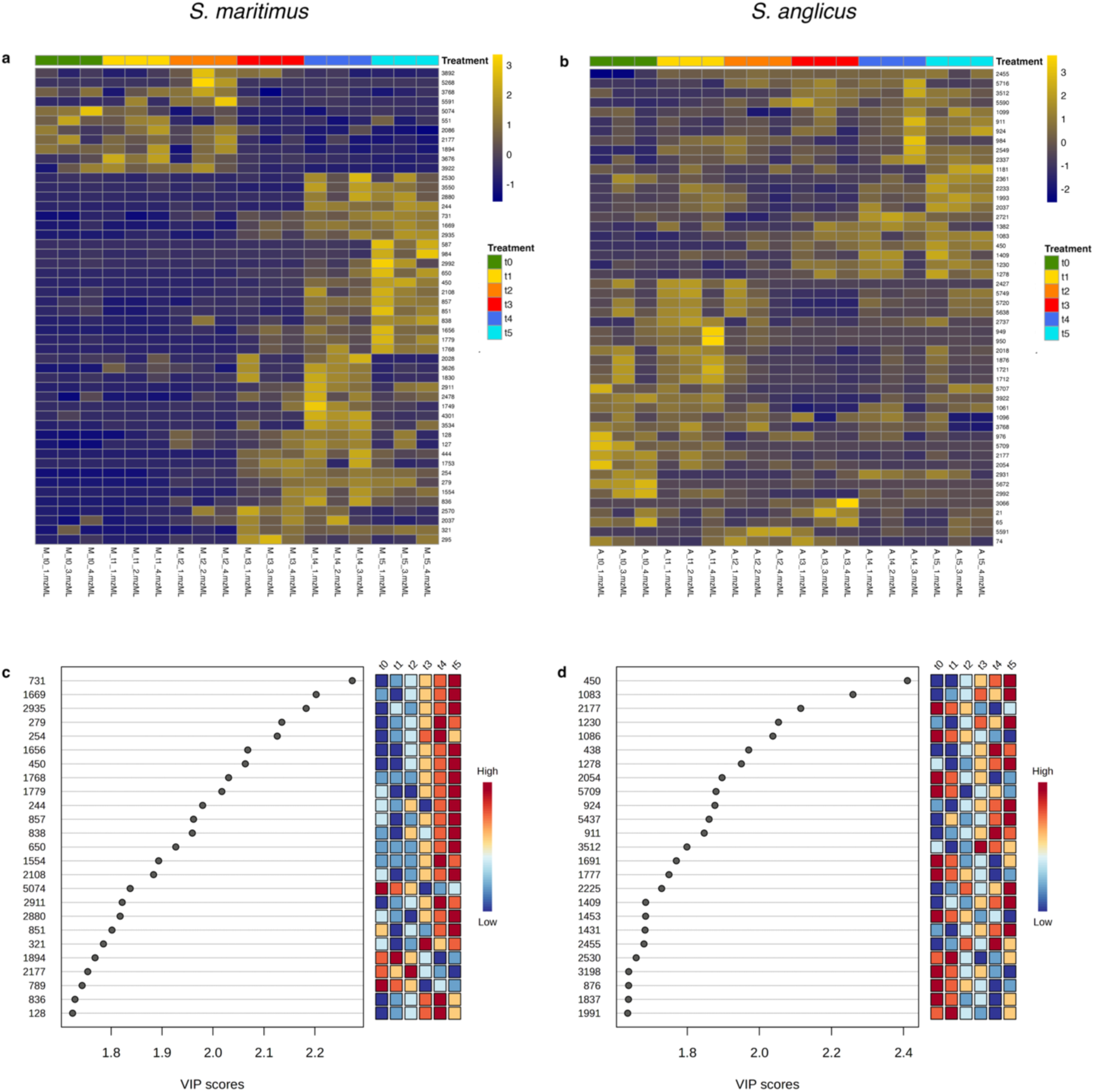
Metabolites variation in the leaves of *S. maritimus* and *S. anglicus*. (**a-b**) Heatmaps showing relative quantification of the 50 most significant features in *S. maritimus* (**a**) and *S. anglicus* (**b**). Colors from blue (low) to yellow (high) indicate the normalized relative abundance values of metabolites. (**c-d**) Partial Least Squares Discriminant Analysis (PLSDA) Variable Importance in Projection (VIP) plots. The y-axis indicates the feature ID, and the x-axis shows the VIP scores (>1.5 indicates a highly influential feature) for the 25 features with the highest scores in *S. maritimus* (**c**) and *S. anglicus* (**d**), along with their quantification across time points. Feature details are provided in Table S22.

## 4. Discussions

Our results demonstrate that *S. maritimus* suffers heat stress more than *S. anglicus*, showing reduced survival, impaired physiological performance, and more pronounced transcriptional changes. In contrast, *S. anglicus* exhibits greater resilience and recovery. In the following sections, we discuss the effects of short-term heat stress (t_0_ *vs*. t_1_), long-term heat stress (t_0_ *vs*. t_3_), and the recovery phase (t_3_ *vs*. t_5_) of both species.

### 4.1 Short-term heat stress

High temperatures greatly affect the photosynthetic apparatus, leading to excess excitation energy (Fortunato *et al*., 2023) and a generation of ROS (Wang *et al*., 2014; Wang *et al*., 2017; Fortunato *et al*., 2023). In our experiment, until t_2_ (36 h of exposure) the heat effects on photosynthetic parameters in both species were limited. Interestingly, both species showed a significant reduction in the chl/car ratio: transient in the NIS (at t_1_) and progressive in the native species (from t_2_ onward). This ratio is known to decrease under stress conditions (Esteban *et al.,* 2015) and suggests an increase of photosynthesis antenna systems with respect to reaction centers (RCs). This is supported by the decrease of the ratio between chl *a* and *b* in *S. anglicus* (indicating less chl *a* than chl *b*) from t_2_ onward. Higher amounts of chl *b* than chl *a* are positively associated with the abundance of light-harvesting complexes (LHCs) (Tanaka & Tanaka, 2011), important in protecting RCs from stress. Gene expression data showed downregulation of genes associated with the BP term “photosynthesis” in both species at the onset of heat stress. Interestingly, only 4 out of 39 genes were shared between species, suggesting differences in early response to heat stress.

Regarding the chloroplast ultrastructure, we observed an increase of PG in both species, as reported in previous works (Arzac *et al*., 2022), already during the short-term stress. Additionally, the photosynthetic membranes started to lose their organization in the native species at t_2_, while the NIS preserved an internal organization, though showing a reduction in the number of grana. This suggests a negative effect on the photosynthetic apparatus and an increase of ROS production, as previously reported in similar experiments (Yin *et al*., 2007; Kumar *et al*., 2012). In the native species, even though the ultrastructure of the chloroplast was clearly affected, there was no concomitant increase of antioxidant enzyme activity, which may have led to an increase in oxidative damage to the organism (Yin *et al*., 2007; Suzuki *et al*., 2013; Raja *et al*., 2020). However, the NIS showed increased activity of several antioxidant defenses at t_2_, particularly GPx, which was significantly upregulated, along with an upward in APx activity, a key enzyme in the hydrogen peroxide scavenging network (Suzuki *et al*., 2013; Raja *et al*., 2020; Fortunato *et al*., 2023). Heat stress and oxidative damage are known to induce protein misfolding, triggering the upregulation of protective protein networks (Sun *et al*., 2002; Trivedi *et al*., 2016; Wang *et al*., 2018; Haq *et al*., 2019; Singh *et al*., 2019). If these responses are insufficient or not properly activated, endoplasmic reticulum stress can occur, potentially leading to programmed cell death (Liu & Howell, 2016; Meyer *et al*., 2019).

Both species exhibited upregulation of genes related to “unfolded protein binding”, indicating activation of molecular chaperone networks known to maintain protein homeostasis and prevent aggregation (Zhao & Liu, 2018; Hu *et al*., 2020). However, the specific upregulation of genes involved in “protein refolding” in *S. anglicus* suggests that only the NIS possesses an actively engaged system for repairing misfolded proteins at this time point.

### 4.2 Effects of long-term heat stress

After 5 days of simulated heatwave, most of the analyzed parameters indicated clear signs of stress. In both species we observed a decrease of F_v_’/F_m_’, with lower values in the native species, indicating more stress (Herrmann *et al*., 1996). The difference between the species was even more pronounced for ϕPSII, which measures the actual efficiency of PSII, that is particularly sensitive to abiotic stress (Sikorski *et al*., 2025). At this stage, it is likely that PSII and the OEC were partially dissociated due to the temperature stress (Wang *et al*., 2018; Hu *et al*., 2020). At the same time, we observed an increase of light energy dissipated as heat (represented by ϕNPQ), especially in the native species. This is a protective mechanism that helps prevent oxidative damage to the photosynthetic apparatus (van Amerongen & Croce, 2025). Moreover, higher values of ϕNPQ in the native species indicate that the organism might not be able to transfer the electrons through the electron transport chain of the chloroplast (Wang *et al*., 2018). On the other hand, *S. anglicus* did not show the same difficulties. Specifically, the dissipation occurs through LHCs (Tang *et al.,* 2007), alleviating the pressure on PSII RCs (Hu *et al.,* 2020). Accordingly, the trend of chl *a*/chl *b* and chl/car ratios decreased in both species, indicating higher concentration of LHCs. Additionally, gene expression data revealed downregulation of genes classified under the BP “pigment metabolic process”, which are involved in chlorophyll biosynthesis, likely reflecting heat-induced degradation of biosynthetic enzymes (Wang *et al*., 2018).

These differences are consistent with their chloroplast ultrastructure. A clear unstacking of grana in the native species was observed, with evident dilation of thylakoid membranes. Conversely, the NIS presented a limited reduction in the grana number, and a regular thylakoid system. Similar effects were also reported in *Brassica campestris* L. (Zou *et al*., 2017; Yamamoto *et al*., 2008).

The loss of photosynthetic apparatus integrity is expected to increase ROS production (Wang *et al*., 2018). In the native species the response was relatively mild, with GPx, CAT, and CCA activities showing an upward trend, although not significantly different from t_0_. APx activity remained unchanged, but gene expression analysis indicated upregulation of APx transcripts, suggesting that the transcriptional response was initiating. In contrast, the NIS exhibited strong APx activity and high CCA, indicating a more pronounced antioxidant response.

High temperature stress also affects carbon metabolism (Sun & Guo, 2016; Hu *et al*., 2020), since the Calvin-Benson cycle depends on photosynthesis products to function. In both species we detected the downregulation of the gene expressing the sedoheptulose-1,7-bisphosphatase (SBPase) (A0A5J9VL29), one of the limiting factors of the cycle (Ding *et al*., 2017; Chen *et al*., 2022; Falcioni *et al*., 2024). This enzyme is also negatively affected by oxidative damage (Ding *et al*., 2017). Interestingly, only in the native species we found the enrichment of the GO term “carbon fixation”, with a clear downregulation of the process. Additionally, we detected an evident change in metabolites content of *S. maritimus*, whilst the NIS did not show the same degree of change. Though many of these metabolites are unknown (annotation level 4), 10 in *S. maritimus* and 8 in *S. anglicus* belong to the superclass “Lipids and lipid-like molecules”, which are known to play an important role in plant stress regulation (*e*.*g*., signaling, initiation of defense reactions, mitigation of stress) (Okazaki & Saito 2014; Hou *et al*., 2016).

High expression of heat shock proteins (HSPs) is generally associated with enhanced stress tolerance (Trivedi *et al*., 2016), although some exceptions have been reported (*e.g.*, HSP90; Haq *et al*., 2019). In the native species, we observed partial downregulation of HSPs and molecular chaperonins, suggesting a less efficient chaperone system compared to the NIS. Consistently, genes associated with the BP term “protein refolding” were also partially downregulated in the native species, whereas in the NIS we detected a clear upregulation of genes enriching the term “response to topologically incorrect proteins”. Furthermore, enrichment of the term “positive regulation of DNA repair” in the native species may indicate the occurrence of DNA damage. Such damage is likely driven by elevated ROS levels (Tutar *et al*., 2017; Haq *et al*., 2019), which appear to be insufficiently scavenged by antioxidant defenses, as discussed above.

### 4.3 Effects of recovery after the simulated heatwave

Following the return to initial temperature conditions, the native species failed to recover the assessed physiological and biochemical parameters with a low survival rate. In contrast, *S. anglicus* showed a concurrent recovery of these parameters to levels comparable to initial conditions. These results indicate that the imposed heat stress exceeded the tolerance threshold of the native species. Consistently, *S. maritimus* exhibited peak activities of APx, GPx, and CAT during the recovery phase, accompanied by an increase in CCA, suggesting a delayed and potentially stress-induced antioxidant response. GO enrichment analysis of the native species at this time point revealed enrichment of terms related to protein binding (*e.g.*, “unfolded protein binding”) and oxidative stress (“response to oxygen-containing compounds”). However, the activation of these response mechanisms likely occurred too late and/or after damage had already become too severe, such as the complete disruption of photosynthetic membranes. In contrast, the NIS showed upregulation of genes encoding proteins involved in the formation and stability of the OEC, indicating active repair of the photosynthetic apparatus. These patterns suggest that while the native species continued to exhibit stress symptoms, the NIS was undergoing recovery. This interpretation is further supported by the upregulation of genes related to “cellular component biogenesis” and “photosynthesis” BP terms, consistent with restoration of photosynthetic function.

## 5. Conclusions

This study highlights the value of integrating ultrastructural, physiological, transcriptomic, and metabolomic approaches to achieve a comprehensive understanding of plant response to stress. By applying these methods to a non-model species, we provide new insights into its biology while expanding available molecular resources. Despite some limitations, including incomplete metabolite identification and constraints in functional annotation of the *de novo* transcriptome, these results lay a solid foundation for future studies and highlight the potential of integrated omics to advance research in non-model systems.

Overall, our results indicate that the NIS responds more rapidly to heat stress, limiting damage to the photosynthetic apparatus and coping more effectively with prolonged stress, compared to *S. maritimus*. Moreover, *S. anglicus* was able to fully recover following stress exposure, restoring photosynthetic efficiency to levels comparable to initial conditions. In contrast, *S. maritimus* failed to recover and exhibited a rapid decline in survival after five days of heat stress. Taken together, these findings indicate that *S. anglicus* is more resilient to heat stress than *S. maritimus*. In the Venice Lagoon, this disparity may contribute to the decline or local disappearance of the native species under increasingly severe stress conditions. This study provides a valuable framework for future research, offering a basis for investigating plant stress responses in plants in complex, multivariable ecosystems.

## Acknowledgments

We thank the Chioggia, Plant Genome Editing and Phenotyping, and Imaging facilities (Department of Biology, University of Padova, Italy) for their support. This work also made use of computational infrastructure funded by the University of Padova Strategic Research Infrastructure Grant 2017 (“CAPRI: Calcolo ad Alte Prestazioni per la Ricerca e l’Innovazione”) and the BioData Hub (Department of Biology, University of Padova, Italy).

## Author contributions

Conceptualization: FD, IM, CDP

Data Curation: FD, PA, RT

Formal Analysis: FD, PA, RT, CF

Funding Acquisition: CDP

Investigation: FD, RT, DC, CS, CF

Methodology: FD, PA, RT, CS, GS, IM, CDP

Project Administration: CDP

Resources: CF, IM, CDP

Supervision: DDB, LA, GS, IM, CDP

Validation: IM, CDP

Visualization: FD, DC, CDP

Writing - Original Draft: FD, PA, RT

Writing - Review & Editing: FD, PA, RT, CS, DC, DDB, CF, LA, GS, IM, CDP

## Data Availability

Raw data for *de novo* transcriptome assembly and differential gene expression analysis are available on the public online repository Sequence Read Archive (SRA) at NCBI, under BioProject accession number PRJNA1269774.

Raw data for metabolomic features identification are available in the MassIVE repository under the ID MSV000101336.

## Supplemental files (titles and legends)

**Fig. S1 Cytofluorometric analysis of nuclear DNA content for species identification.** A single major peak (representing the G1 phase of the cell cycle) with a small additional peak (representing nuclei in the G2 phase) indicates a uniform population of nuclei in terms of genome size. Two major peaks suggest the presence of two distinct nuclei populations with different genome sizes. Panels **a** and **b** show the nuclei size distribution extracted from a pool of leaves from four plants, hypothesized to be *S. maritimus* (Pool M, M1+M2+M3+M4) (**a**) and *S. anglicus* (Pool A, A1+A2+A3+A4) (**b**), from each pot. Panels **c**, **d**, **e**, and **f** display the genome size of the nuclei extracted from pairs of *S. maritimus* and *S. anglicus* leaves (the same used for Pool M and Pool A), indicating the presence of nuclei with different genome sizes, thus different species (*S. maritimus* and *S. anglicus*).

**Fig. S2 Temperature variation in the growth chamber during the experimental period**. Variation of Temperature (°C) in the growth chamber during the experimental period. Yellow indicates day-time conditions (lights on) and blue indicates night-time conditions (lights off).

**Fig. S3 Changes in specific carotenoid content before, during, and after heatwave stress.** Quantification of Neoxanthin (**a-b**), Violaxanthin (**c-d**), Lutein (**e-f**), Zeaxanthin (**g-h**), and β-Carotene (**i-j**) in *S. maritimus* (**a, c, e, g, i**) and *S. anglicus* (**b, d, f, h, j)** from t_0_ to t_5_. The results (n=3) are expressed as mean ± SD. One-way ANOVA with Tukey’s post-hoc test was carried out to determine significant differences; results are reported in Tables S5 and S6. No statistically significant differences were found.

**Fig. S4 Number of differentially expressed genes (DEGs) across all comparisons.** DEG counts for *S. maritimus* (green) and *S. anglicus* (orange) under short-term heat stress (t_0_ *vs.* t_1_), long-term heat stress (t_0_ *vs.* t_3_), recovery phase (t_3_ *vs.* t_5_) and comparison with initial conditions (t_0_ *vs.* t_5_).

**Fig. S5 GO analysis of differentially expressed genes in *S. anglicus* at the end of the experiment (t_0_ *vs.* t_5_).** Significantly enriched Biological Processes (BP, **a**), Molecular Functions (MF, **b**), and Cellular Components (CC, **c**) in *S*. *anglicus* are shown for the comparison between the start and end of the experiment (t_0_ *vs.* t_5_). Circle size is proportional to the number of DEGs and color (blue-low, yellow-high) is proportional to the statistical significance, expressed as the negative logarithm of the False Discovery Rate (-log10(FDR)). Complete DEGs lists and enriched GO terms are provided in Tables S14 and S21.

**Fig. S6 Enriched Molecular Function (MF) terms in the leaves of *S. maritimus* and *S. anglicus*.** Significantly enriched MF GO terms are shown for the different comparisons: (**a–b**) short-term heat stress (t_0_ *vs.* t_1_); (**c–d**) long-term heat stress (t_0_ *vs.* t_3_); (**e–f**) recovery phase (t_3_ *vs.* t_5_). Circle size is proportional to the number of DEGs and color (blue-low, yellow-high) is proportional to the statistical significance, expressed as the negative logarithm of the False Discovery Rate (-log10(FDR)). Complete DEGs lists and enriched GO terms for both species are provided in Tables S7, S8, S9, and S15, S16, S17 (*S. maritimus*) and S11, S12, S13 and S18, S19, S20 (*S. anglicus*)

**Fig. S7 Expression profiles of genes associated with the GO term “unfolded protein binding” in *S. maritimus* and *S. anglicus*.** Heatmaps show the absolute expression values of genes in *S. maritimus* (**a, c, e**) and *S. anglicus* (**b, d, f**) during short-term heat stress (t_0_ *vs*. t_1_; **a, b**), long-term heat stress (t_0_ *vs*. t_3_; **c, d**), and the recovery phase (t_3_ *vs*. t_5_; **e, f**) associated with the GO term “unfolded protein binding” (GO:0006986). The UniProt identifier is reported on each row. The color of the cell is proportional to the expression value (blue-low, yellow-high), represented as log2.

**Fig. S8 Expression profiles of genes associated with GO terms “pigment metabolic process” and “carbon fixation” in *S. maritimus*.** Heatmaps show the absolute expression values of genes associated with “pigment metabolic process” (GO:0042440) (**a**) and “carbon fixation” (GO:0015977) (**b**) in *S. maritimus* during long-term heat stress (t_0_ *vs*. t_3_). The UniProt identifier is reported on each row. The color of the cell is proportional to the expression value (blue-low, yellow-high), represented as log2.

**Fig. S9 Enriched Cellular Component (CC) terms in the leaves of *S. maritimus* and *S. anglicus*.** Significantly enriched CC terms are shown for the different comparisons: short-term heat stress (t_0_ *vs*. t_1_; **a–b**), long-term heat stress (t_0_ *vs*. t_3_; **c–d**), and recovery phase (t_3_ *vs*. t_5_; **e–f**). Circle size is proportional to the number of DEGs and color (blue-low, yellow-high) is proportional to the statistical significance, expressed as the negative logarithm of the False Discovery Rate (-log10(FDR)). Complete DEGs lists and enriched GO terms for both species are provided in Tables S7, S8, S9 and S15, S16, S117 (*S. maritimus*) and S11, S12, S13 and S18, S19, S20 (*S. anglicus*).

**Table S1 List of primers for species identification.** The acronym of the marker used (chloroplast *trnL-trnF*, nuclear ITS), the sequence (5’-3’) and the reference are reported.

**Table S2 Species identification combining gene marker analysis and genome size evaluation.** Alignment results of *trnL-trnF* and ITS sequences for the sampled organisms are shown. The number of peaks detected by cytofluorometry is also reported, with two peaks indicating nuclei with different genome sizes. By combining genome size data with sequence alignment results, each organism was successfully identified.

**Table S3 Parameters used for feature extraction using MzMine 3.**

**Table S4 Parameters used for spectral library search in GNPS.**

**Table S5 Summary of statistical analysis for *S. maritimus***. Results of Tukey’s post hoc test for significant findings from physiological and biochemical assays conducted on *S. maritimus*.

**Table S6 Summary of statistical analysis *S. anglicus*.** Results of Tukey’s post hoc test for significant findings from physiological and biochemical assays conducted on *S. anglicus*.

**Table S7 Differentially expressed genes in *S. maritimus* between t_0_ and t_1_.** List of differentially expressed genes (3’470; FDR <0.01) in leaves of *S. maritimus* after short-term heat stress (t_0_ vs. t_1_).

**Table S8 Differentially expressed genes in *S. maritimus* between t_0_ and t_3_.** List of differentially expressed genes (11’049; FDR <0.01) in leaves of *S. maritimus* after long-term heat stress (t_0_ *vs.* t_3_).

**Table S9 Differentially expressed genes in *S. maritimus* between t_3_ and t_5_.** List of differentially expressed genes (7’744; FDR <0.01) in leaves of *S. maritimus* after the recovery phase (t_3_ *vs.* t_5_).

**Table S10 Differentially expressed genes in *S. maritimus* between t_0_ and t_5_**. List of differentially expressed genes (171; FDR <0.01) in leaves of *S. maritimus* after the end of the experiment (t_0_ *vs.* t_5_).

**Table S11 Differentially expressed genes in *S. anglicus* between t_0_ and t_1_.** List of differentially expressed genes (1’816; FDR <0.01) in leaves of *S. anglicus* after short-term heat stress (t_0_ *vs.* t_1_).

**Table S12 Differentially expressed genes in *S. anglicus* between t_0_ and t_3_.** List of differentially expressed genes (2’034; FDR <0.01) in leaves of *S. anglicus* after long-term heat stress (t_0_ *vs.* t_3_).

**Table S13 Differentially expressed genes in *S. anglicus* between t_3_ and t_5_.** List of differentially expressed genes (6’005; FDR <0.01) in leaves of *S. anglicus* after the recovery phase (t_3_ *vs.* t_5_).

**Table S14 Differentially expressed genes in *S. anglicus* between t_0_ and t_5_.** List of differentially expressed genes (1’594; FDR <0.01) in leaves of *S. anglicus* after the end of the experiment (t_0_ *vs.* t_5_).

**Table S15 Enriched GO terms in *S. maritimus* between t_0_ and t_1_.** List of enriched GO terms (FDR <0.05) in leaves of *S. maritimus* after short-term heat stress (t_0_ *vs.* t_1_).

**Table S16 Enriched GO terms in *S. maritimus* between t_0_ and t_3_.** List of enriched GO terms (FDR <0.05) in leaves of *S. maritimus* after long-term heat stress (t_0_ *vs.* t_3_).

**Table S17 Enriched GO terms in *S. maritimus* between t_3_ and t_5_.** List of enriched GO terms (FDR <0.05) in leaves of *S. maritimus* after the recovery phase (t_3_ *vs.* t_5_).

**Table S18 Enriched GO terms in *S. anglicus* between t_0_ and t_1_.** List of enriched GO terms (FDR <0.05) in leaves of *S. anglicus* after short-term heat stress (t_0_ *vs.* t_1_).

**Table S19 Enriched GO terms in *S. anglicus* between t_0_ and t_3_.** List of enriched GO terms (FDR <0.05) in leaves of *S. anglicus* after long-term heat stress (t_0_ *vs.* t_3_).

**Table S20 Enriched GO terms in *S. anglicus* between t_3_ and t_5_.** List of enriched GO terms (FDR <0.05) in leaves of *S. anglicus* after the recovery phase (t_3_ *vs.* t_5_).

**Table S21 Enriched GO terms in *S. anglicus* between t_0_ and t_5_.** List of enriched GO terms (FDR <0.05) in leaves of *S. anglicus* after the end of the experiment (t_0_ *vs.* t_5_).

**Table S22 Detected metabolic features in *S. maritimus* and *S. anglicus*.** Metabolites detected by Ultra-Performance Liquid Chromatography (UPLC) in leaves of *S. maritimus* and *S. anglicus*. Reported information includes Feature ID, mass-to-charge ratio (mz), retention time (rt), adduct type, putative molecular formula, annotation level, annotated name for features annotated with level 2, and compound classification.

