## Supplementary Figures and Tables (1 to 4) for "Grass wars: how native and non-indigenous *Sporobolus* battle heatwaves in salt marshes"

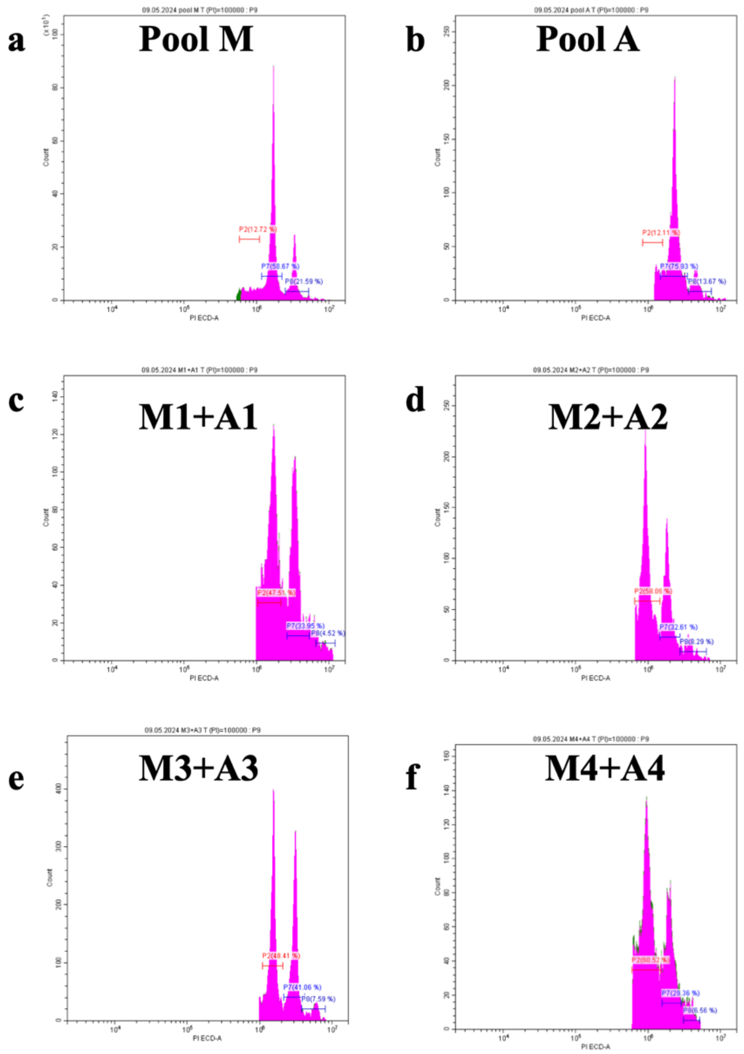


**Fig. S1 Cytofluorometric analysis of nuclear DNA content for species identification.** A single major peak (representing the G1 phase of the cell cycle) with a small additional peak (representing nuclei in the G2 phase) indicates a uniform population of nuclei in terms of genome size. Two major peaks suggest the presence of two distinct nuclei populations with different genome sizes. Panels **a** and **b** show the nuclei size distribution extracted from a pool of leaves from four plants, hypothesized to be *S. maritimus* (Pool M, M1+M2+M3+M4) (**a**) and *S. anglicus* (Pool A, A1+A2+A3+A4) (**b**), from each pot. Panels **c**, **d**, **e**, and **f** display the genome size of the nuclei extracted from pairs of *S. maritimus* and *S. anglicus* leaves (the same used for Pool M and Pool A), indicating the presence of nuclei with different genome sizes, thus different species (*S. maritimus* and *S. anglicus*)

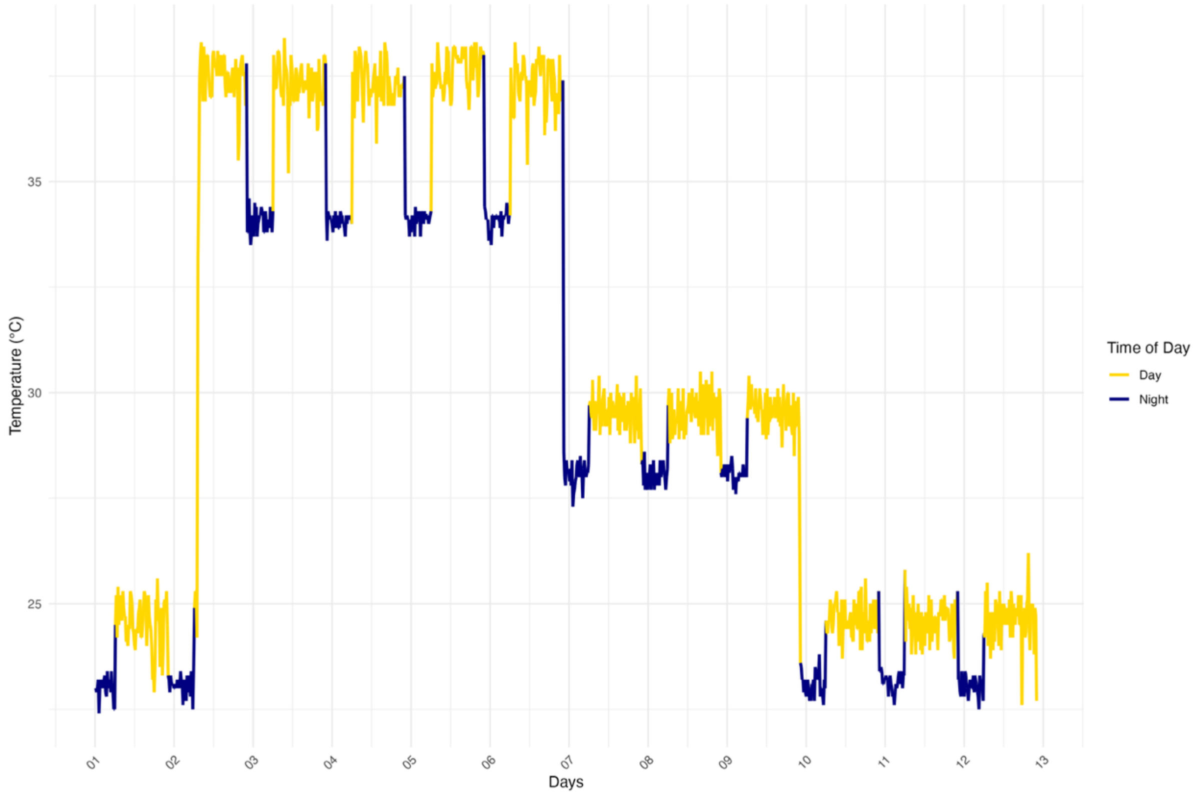


**Fig. S2 Temperature variation in the growth chamber during the experimental period**. Variation of Temperature (°C) in the growth chamber during the experimental period. Yellow indicates day-time conditions (lights on) and blue indicates night-time conditions (lights off).

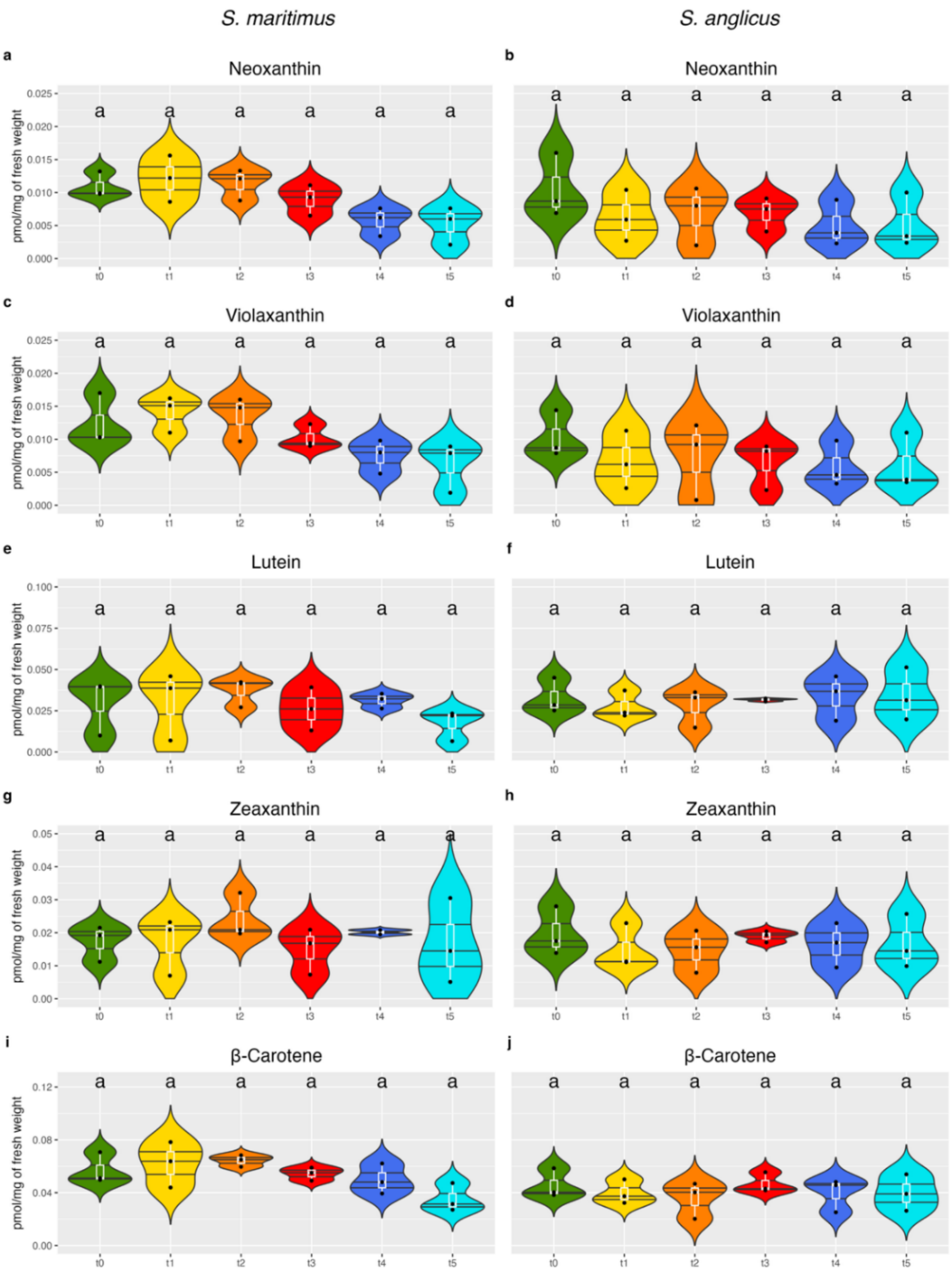


**Fig. S3 Changes in specific carotenoid content before, during, and after heatwave stress.** Quantification of Neoxanthin (**a-b**), Violaxanthin (**c-d**), Lutein (**e-f**), Zeaxanthin (**g-h**), and β-Carotene (**i-j**) in *S. maritimus* (**a, c, e, g, i**) and *S. anglicus* (**b, d, f, h, j)** from $t_0$ to $t_5$. The results (n=3) are expressed as mean ± SD. One-way ANOVA with Tukey's post-hoc test was carried out to determine significant differences; results are reported in Tables S5 and S6. No statistically significant differences were found

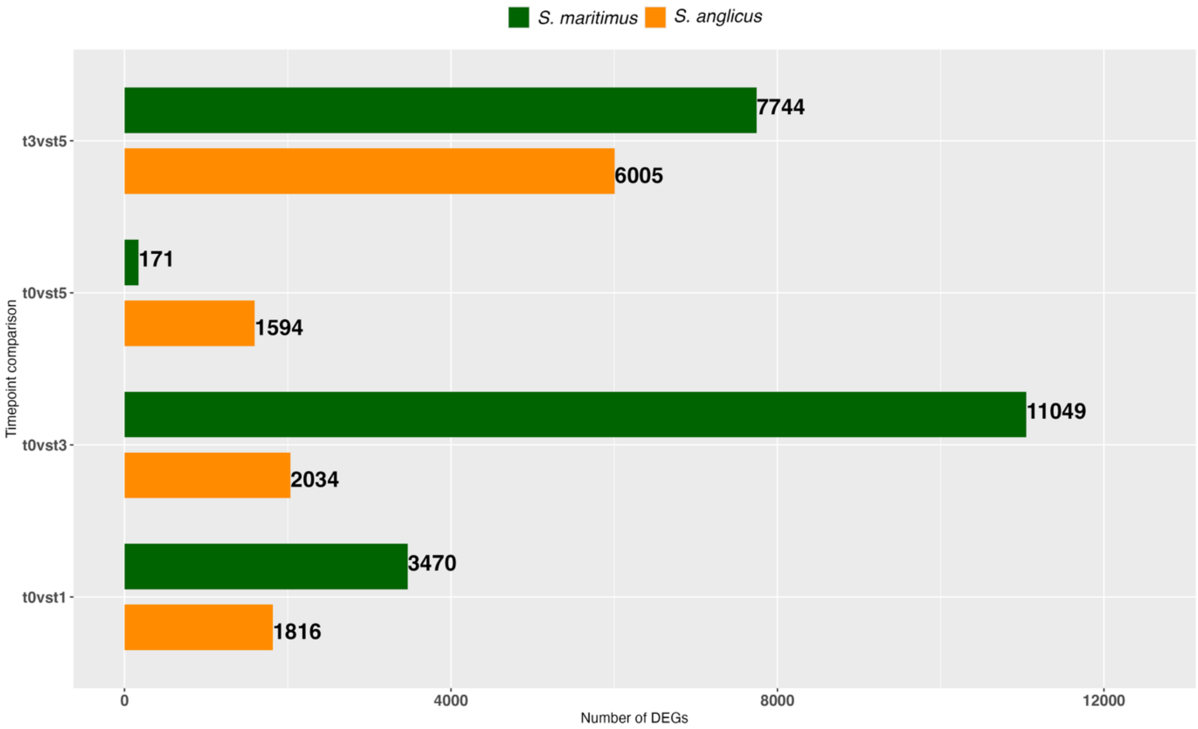


**Fig. S4 Number of differentially expressed genes (DEGs) across all comparisons.** DEG counts for *S. maritimus* (green) and *S. anglicus* (orange) under short-term heat stress ($t_0$ *vs.* $t_1$), long-term heat stress ($t_0$ *vs.* $t_3$), recovery phase ($t_3$ *vs.* $t_5$) and comparison with initial conditions ($t_0$ *vs.* $t_5$).

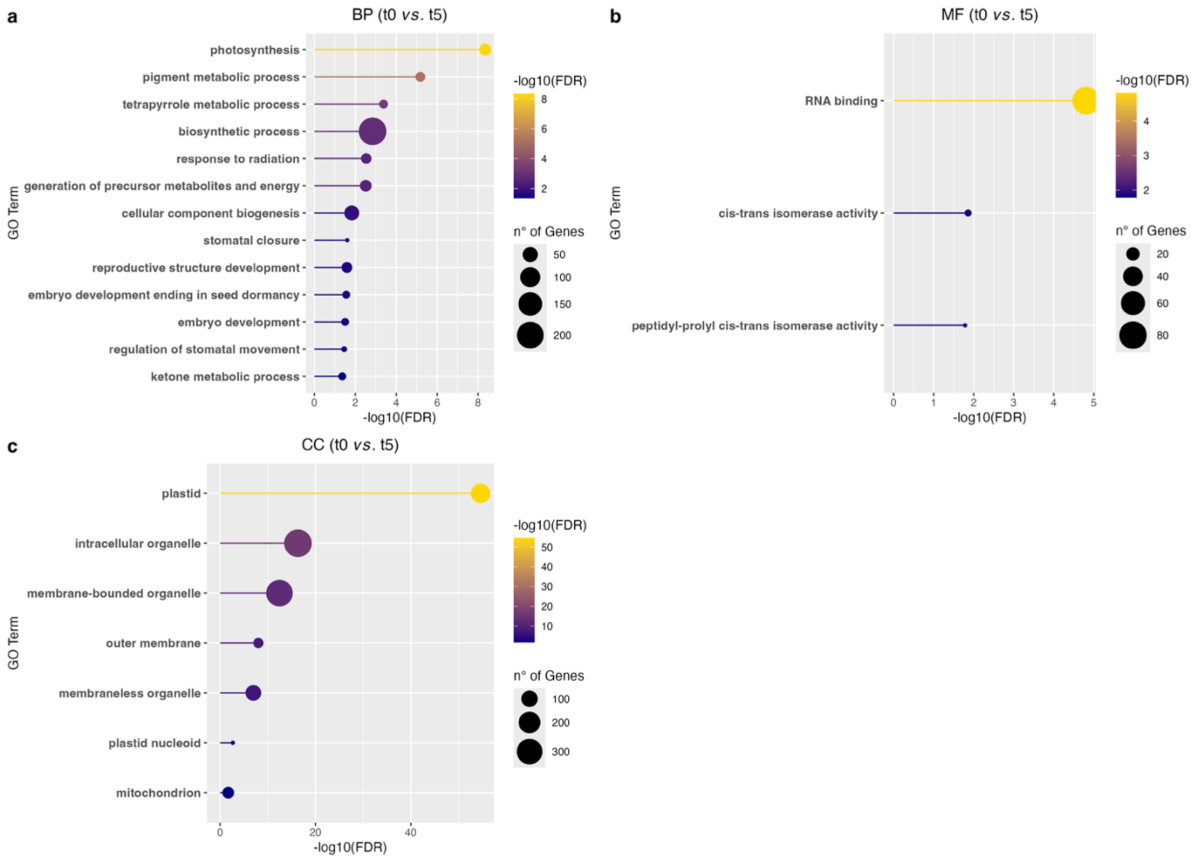


**Fig. S5 GO analysis of differentially expressed genes in *S. anglicus* at the end of the experiment ($t_0$ *vs.* $t_5$).** Significantly enriched Biological Processes (BP, **a**), Molecular Functions (MF, **b**), and Cellular Components (CC, **c**) in *S. anglicus* are shown for the comparison between the start and end of the experiment ($t_0$ *vs.* $t_5$). Circle size is proportional to the number of DEGs and color (blue-low, yellow-high) is proportional to the statistical significance, expressed as the negative logarithm of the False Discovery Rate (-log10(FDR)). Complete DEGs lists and enriched GO terms are provided in Tables S14 and S21.

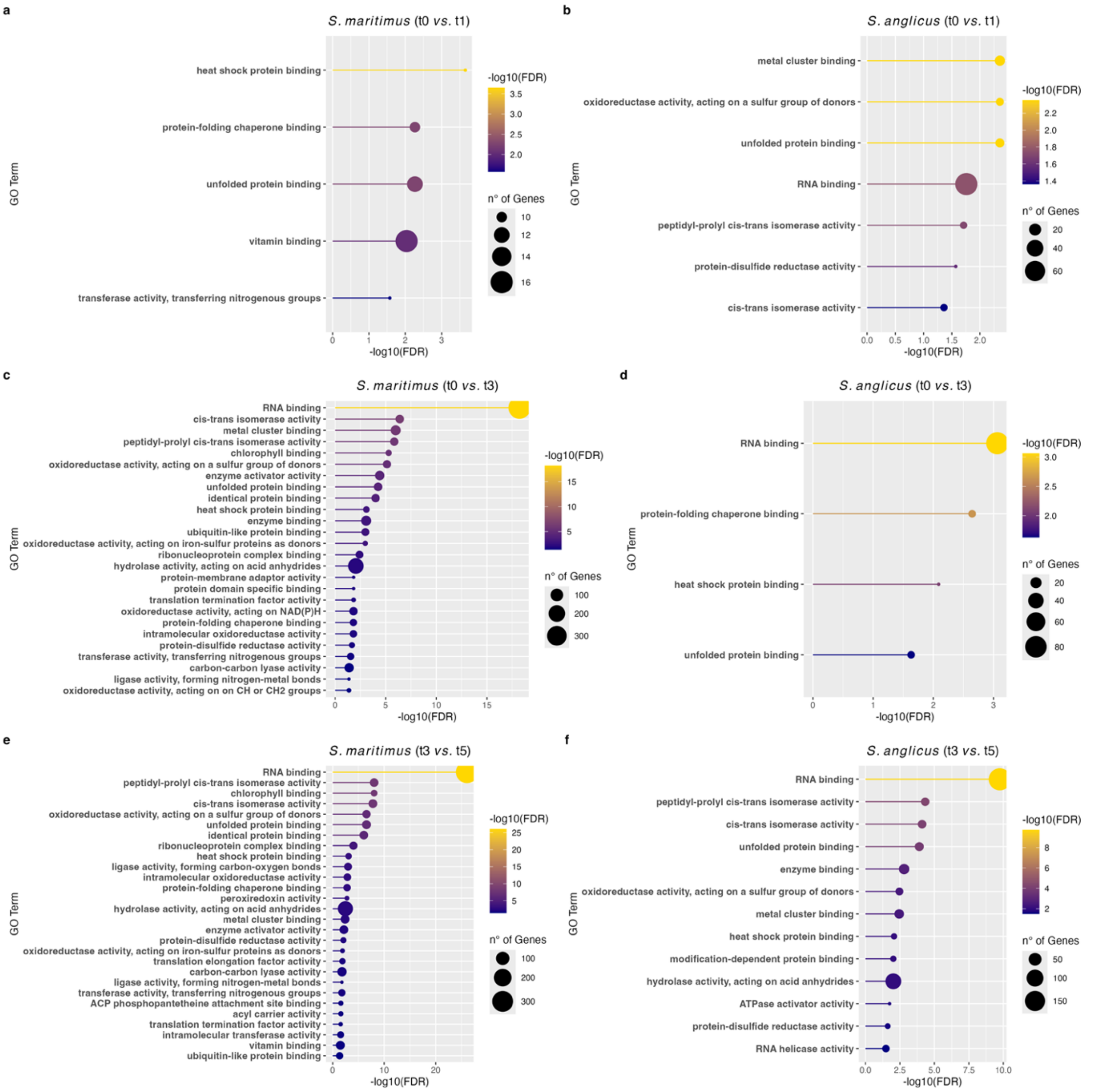


**Fig. S6 Enriched Molecular Function (MF) terms in the leaves of *S. maritimus* and *S. anglicus*.** Significantly enriched MF GO terms are shown for the different comparisons: (**a–b**) short-term heat stress ($t_0$ *vs.* $t_1$); (**c–d**) long-term heat stress ($t_0$ *vs.* $t_3$); (**e–f**) recovery phase ($t_3$ *vs.* $t_5$). Circle size is proportional to the number of DEGs and color (blue-low, yellow-high) is proportional to the statistical significance, expressed as the negative logarithm of the False Discovery Rate (-log10(FDR)). Complete DEGs lists and enriched GO terms for both species are provided in Tables S7, S8, S9, and S15, S16, S17 (*S. maritimus*) and S11, S12, S13 and S18, S19, S20 (*S. anglicus*)

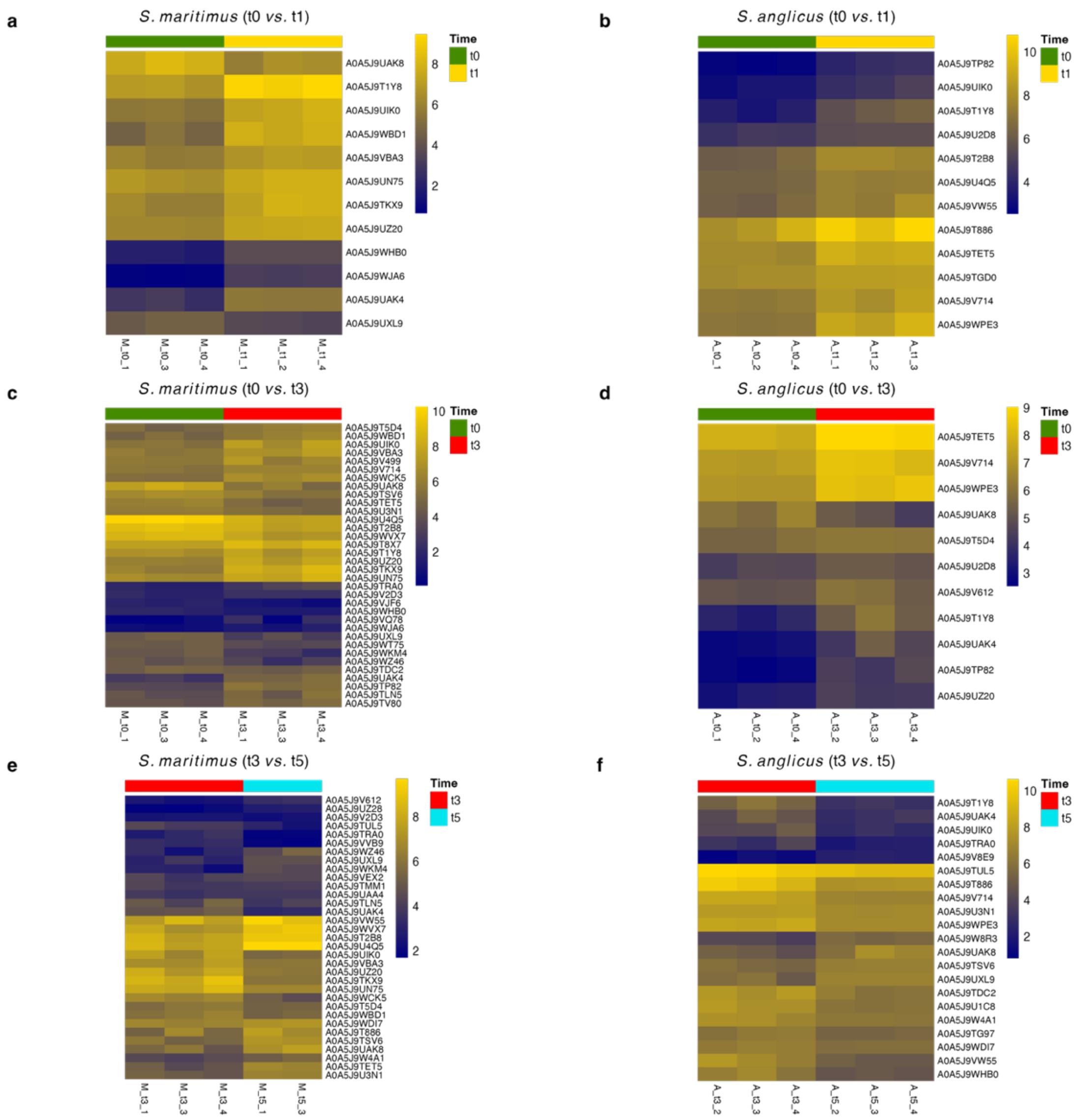


**Fig. S7 Expression profiles of genes associated with the GO term "unfolded protein binding" in *S. maritimus* and *S. anglicus*.** Heatmaps show the absolute expression values of genes in *S. maritimus* (**a, c, e**) and *S. anglicus* (**b, d, f**) during short-term heat stress ($t_0$ *vs*. $t_1$; **a, b**), long-term heat stress ($t_0$ *vs*. $t_3$; **c, d**), and the recovery phase ($t_3$ *vs*. $t_5$; **e, f**) associated with the GO term "unfolded protein binding" (GO:0006986). The UniProt identifier is reported on each row. The color of the cell is proportional to the expression value (blue-low, yellow-high), represented as log2.

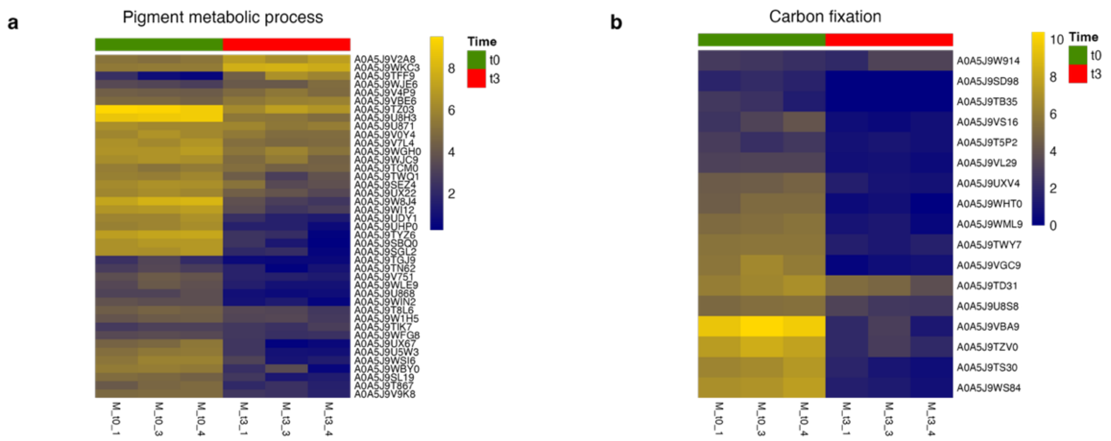


**Fig. S8 Expression profiles of genes associated with GO terms "pigment metabolic process" and "carbon fixation" in *S. maritimus*.** Heatmaps show the absolute expression values of genes associated with "pigment metabolic process" (GO:0042440) (**a**) and "carbon fixation" (GO:0015977) (**b**) in *S. maritimus* during long-term heat stress ($t_0$ *vs*. $t_3$). The UniProt identifier is reported on each row. The color of the cell is proportional to the expression value (blue-low, yellow-high), represented as log2.

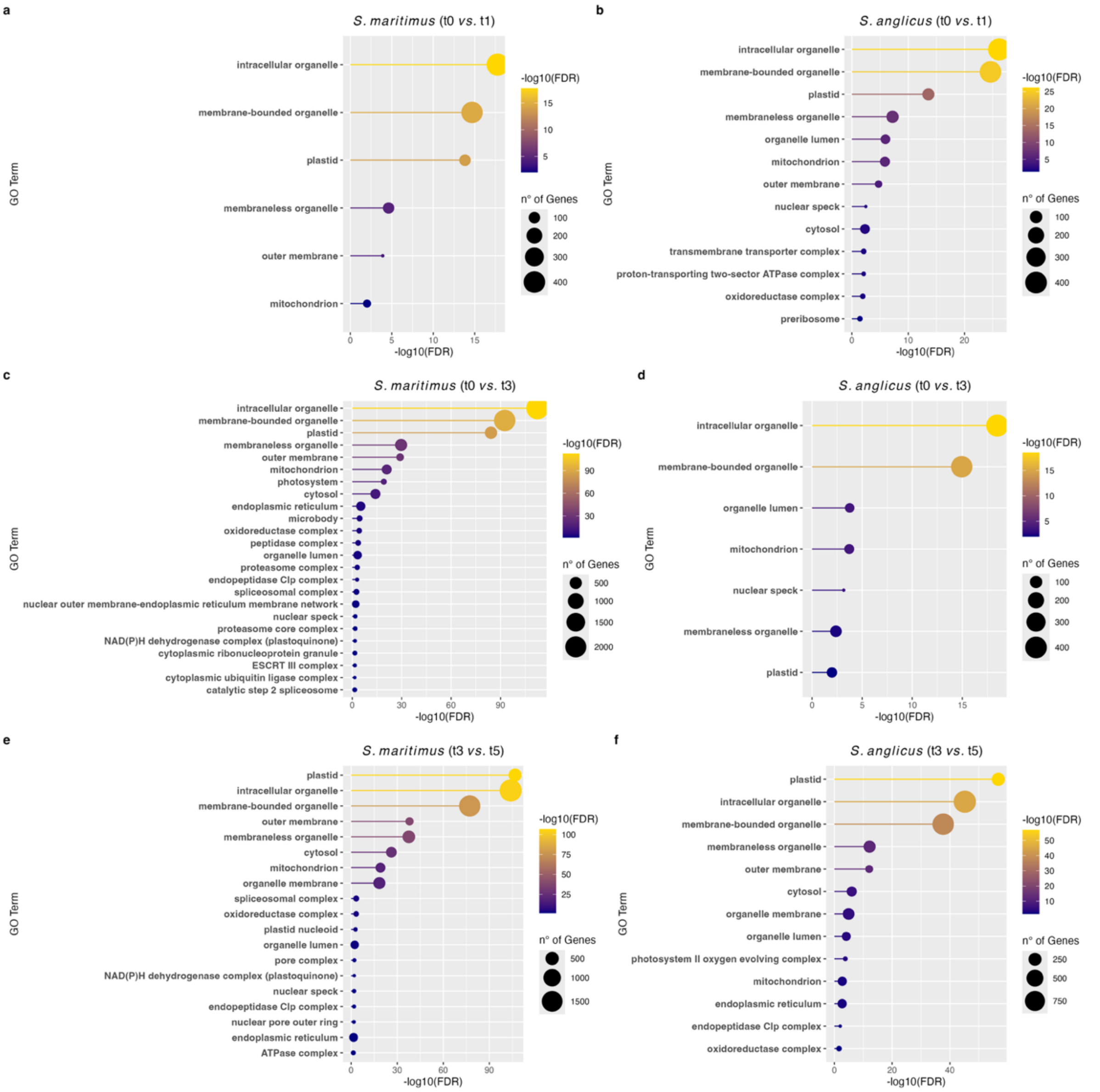


**Fig. S9 Enriched Cellular Component (CC) terms in the leaves of *S. maritimus* and *S. anglicus*.** Significantly enriched CC terms are shown for the different comparisons: short-term heat stress ($t_0$ *vs*. $t_1$; **a–b**), long-term heat stress ($t_0$ *vs*. $t_3$; **c–d**), and recovery phase ($t_3$ *vs*. $t_5$; **e–f**). Circle size is proportional to the number of DEGs and color (blue-low, yellow-high) is proportional to the statistical significance, expressed as the negative logarithm of the False Discovery Rate (-log10(FDR)). Complete DEGs lists and enriched GO terms for both species are provided in Tables S7, S8, S9 and S15, S16, S117 (*S. maritimus*) and S11, S12, S13 and S18, S19, S20 (*S. anglicus*).

**Table S1 List of primers for species identification.** The acronym of the marker used (chloroplast *trnL-trnF*, nuclear ITS), the sequence (5'-3') and the reference are reported.

| Marker | Primer | Sequence (5' - 3') | Reference |
|---|---|---|---|
| *trnL-trnF* | F | ATAGATCCTGACATAGCAAACGAT | This study |
| | R | TCATCCTAGTAGAGTATTTGCATCC | This study |
| ITS | F | CATCCATGGCATCGGGTGGC | Muhammad & Ki, 2022 |
| | R | GGGGCAATGTCACTGAATTCAG | Muhammad & Ki, 2022 |

**Table S2 Species identification combining gene marker analysis and genome size evaluation.** Alignment results of *trnL-trnF* and ITS sequences for the sampled organisms are shown. The number of peaks detected by cytofluorometry is also reported, with two peaks indicating nuclei with different genome sizes. By combining genome size data with sequence alignment results, each organism was successfully identified.

| **Sample** | **Chloroplast marker (*trnL-trnF*)** | **Nuclear marker (*ITS*)** | **Peaks at the cytofluorometer with a morphologically different organism** | **Identification** |
|---|---|---|---|---|
| M1 | *S. maritimus* | *S. maritimus* | 2 | *S. maritimus* |
| M2 | *S. maritimus* | *S. maritimus* | 2 | *S. maritimus* |
| M3 | *S. maritimus* | *S. maritimus* | 2 | *S. maritimus* |
| M4 | *S. maritimus* | *S. maritimus* | 2 | *S. maritimus* |
| A1 | *S. anglicus* / *S. alterniflorus* | *S. maritimus* | 2 | *S. anglicus* |
| A2 | *S. anglicus* / *S. alterniflorus* | *S. maritimus* | 2 | *S. anglicus* |
| A3 | *S. anglicus* / *S. alterniflorus* | *S. maritimus* | 2 | *S. anglicus* |
| A4 | *S. anglicus* / *S. alterniflorus* | *S. maritimus* | 2 | *S. anglicus* |

**Table S3 Parameters used for feature extraction using MzMine 3.**

| | | |
|---|---|---|
| **Mass detection** | | |
| Scans | MS level | 1 |
| | Polarity | Any |
| | Spectrum | Any |
| Mass detector | Centroid | |
| | Noise Level | $5.00 * 10^3$ |
| Scans | MS level | 2 |
| | Polarity | Any |
| | Spectrum | Any |
| Mass detector | Centroid | |
| | Noise Level | $2.00 * 10^2$ |
| **ADAP Chromatogram Builder** | | |
| Scans | MS level | 1 |
| | Polarity | Any |
| | Spectrum type | Any |
| Min group size in # of scans | | 5 |
| Group intensity threshold | | $1.50 * 10^4$ |
| Min highest intensity | | $5.00 * 10^4$ |
| Scan to scan accuracy | | 0.005 m/z to 10 ppm |
| **Local Minimum Feature Resolver** | | |
| MS/MS scan pairing | Retention time tolerance | 0.05 |
| | MS1 to MS2 precursor tolerance | 0.01 m/z to 10 ppm |
| Chromatographic threshold | | 95% |
| Minimum search range RT/Mobility (absolute) | | 0.25 |
| Minimum absolute height | | $5.00 * 10^4$ |
| Min ratio of peak top/edge | | 1.90 |
| Peak duration range (min/mobility) | | 0.00-2.00 |
| Min # of data points | | 7 |
| **13C isotope filter** | | |
| m/z tolerance | | 0.001 m/z or 3 ppm |
| Retention time tolerance | | 0.02 |
| Monotonic shape | | ✓ |
| Maximum charge | | 2 |
| Representative isotope | | Most intense |
| Never remove feature with MS2 | | ✓ |
| **Isotopic peaks finder** | | |
| Chemical elements | | C, N, O, Cl, Br |
| m/z tolerance | | 0.001 m/z or 5 ppm |
| Maximum charge of isotope m/z | | 2 |
| Search in scans | | Single most intense |
| **Join aligner** | | |
| m/z tolerance | | 0.001 m/z or 5 ppm |
| Weight for m/z | | 3 |
| Retention time tolerance | | 0.06 |
| Weight for RT | | 1 |
| Mobility weight | | 1 |
| **Peak list rows filter** | | |
| Minimum features in a row (abs or %) | ✓ | 4 |
| Validate C13 isotope pattern | m/z tolerance | 0.0005 m/z or 5 ppm |
| | Max charge | 2 |
| | Estimate minimum carbon | ✓ |

| | | |
|---|---|---|
| | Remove if C13 | ✓ |
| | Exclude isotope | O |
| Keep or remove rows | Keep rows that match all criteria | |
| Never remove features with MS2 | ✓ | |

**Table S4 Parameters used for spectral library search in GNPS.**

| Search Options | |
|---|---|
| Precursor Ion Mass Tolerance | ± 0.025 Da |
| Fragment Ion Mass Tolerance | ± 0.02 Da |
| Min Matched Peaks | 4 |
| Score Threshold | 0.75 (cosine score) |
| **Advanced Search Options** | |
| Library Class | Bronze |
| Top Hits Per Spectrum | 1 |
| Search Analogs | Don't search |
| **Advanced Filtering Options** | |
| Filter StdDev Intensity | 0 |
| Filter Precursor Window | Filter |
| Filter peaks in 50Da Window | Filter |
| Filter SNR Intensity | 0 |
| Filter Library | Filter library |
| Min Peak Int | 0 |
